# Impaired value-based decision-making in Parkinson’s Disease Apathy

**DOI:** 10.1101/2023.07.27.550708

**Authors:** William Gilmour, Graeme Mackenzie, Mathias Feile, Louise Tayler-Grint, Szabolcs Suveges, Jennifer A Macfarlane, Angus D Macleod, Vicky Marshall, Iris Q Grunwald, J Douglas Steele, Tom Gilbertson

## Abstract

Apathy is a common and disabling complication of Parkinson’s disease characterised by reduced goal-directed behaviour. Several studies have reported dysfunction within pre-frontal cortical regions and projections from brainstem nuclei whose neuromodulators include dopamine, serotonin and noradrenaline. Work in animal and human neuroscience have confirmed contributions of these neuromodulators on aspects of motivated decision making. Specifically, non-dopaminergic neuromodulators, influence decisions to explore alternative courses of action or persist in an existing strategy to achieve a rewarding goal.

Building upon this work, we hypothesised that Apathy in Parkinson’s disease should be associated with a failure to adequately monitor and make adaptive choices when the rewarding outcome of decisions are uncertain. Using a 4-armed restless bandit reinforcement learning task, we studied decision making in 75 volunteers; 53 patients with Parkinson’s disease, with and without clinical apathy, and 22 age matched healthy controls. Patients with Apathy exhibited impaired ability to choose the highest value bandit. Task performance predicted an individual patient’s apathy severity measured using the Lille Apathy Rating scale (R = -0.46, p<0.001). Computational modelling of the patient’s choices confirmed the apathy group made decisions that that were indifferent to the learnt value of the options, consistent with previous reports of reward insensitivity. Further analysis demonstrated a shift away from exploiting the highest value option and a reduction in perseveration which also correlated with apathy scores (R = -0.5, p<0.001).

We went on to acquire fMRI in 59 volunteers; a group of 19 patients with and 20 without apathy and 20 age matched controls performing the restless bandit task. Analysis of the fMRI signal at the point of reward feedback confirmed diminished signal within ventromedial prefrontal cortex in Parkinson’s disease, which was more marked in Apathy, but not predictive of their individual Apathy severity. Using a model-based categorisation of choice type, decisions to explore lower value bandits in the apathy group activated pre-frontal cortex to a similar degree to the age-matched controls. In contrast, Parkinson’s patients *without* apathy demonstrated significantly increased activation across a distributed thalamo-cortical network. Enhanced activity in the thalamus predicted individual apathy severity across both patient groups and exhibited functional connectivity with dorsal anterior cingulate cortex and anterior insula.

Given that task performance in patients without apathy was no different to the age-matched controls, we interpret the recruitment of this network as a possible compensatory mechanism, which compensates against symptomatic manifestation of apathy in Parkinson’s disease.

## Introduction

Apathy is a debilitating and poorly understood syndrome characterised by a reduction in goal-directed behaviour. In neurodegenerative conditions including Parkinson’s Disease (PD) it is estimated to affect 30-70% of patients ^1–5^. In contrast to the defining motor symptoms of this condition, apathy is a greater predictor of poor quality of life ^6^. Apathy increases the likelihood of developing Dementia ^7^ and presents a clinical challenge, due to its resistance to dopamine replacement or any other treatments ^8^. Importantly, apathy is understood to be a distinct process and a direct consequence of neurodegeneration, independent from co-morbid mood disorder or a secondary consequence of physical disability of the disease ^2, 9^. Although the burden of apathy on patients with neurodegenerative disease is increasingly recognised clinically ^10^, limited progress has been made in understanding the neural circuit mechanisms ^11^. Understanding these is crucial to developing novel treatments, as different dysfunctional neural process are likely to contribute to the manifestation of apathy in different clinical populations ^12^.

Neuroimaging studies of PD patients with apathy have consistently identified either structural or functional imaging abnormalities within pre-frontal cortical circuits and their reciprocally connected sub-cortical nuclei of the basal ganglia ^13^. These abnormalities include the orbitofrontal cortex (OFC) ^14, 15^ ventromedial prefrontal cortex ^16–18^, anterior cingulate cortex (ACC) ^15, 18^ caudate, and ventral striatum ^13, 15, 16^. Abnormalities of brain volume and functional activation have also been localised to the same areas in patients with apathy with different neurodegenerative conditions ^19, 20^ supporting a transdiagnostic anatomical basis.

The function of this fronto-striatal circuit in decision making has been refined by decades of cognitive neuroscience research and include discrete contributions to evaluation of effort costs, choice arbitration, and the encoding of the value of actions and sensory stimuli ^21–24^. On the basis of these functions and reviewing neuroimaging studies of apathy, Le Heron *et. al*., (2017) ^19^ proposed that apathy could arise from dysfunction within a fronto-striatal circuit that mediates any of three key elements of motivated behaviour (i) deciding whether to act; (ii) persisting with an action; and (iii) learning, through outcome monitoring, whether a behaviour was worth performing.

In support of a higher threshold for (i) deciding whether to act; when faced with the decision to exert effort for a monetary reward, apathetic patients tend to reject more offers compared to non-apathetic counterparts ^25^. This behaviour is not due to a heightened sensitivity to effort costs, but rather points to a diminished incentivization by rewarding outcomes, a characteristic feature of apathy ^25^. This interpretation aligns with the observations of reward insensitivity as a general feature of apathy in PD, corroborated by diminished pupillary response ^26^, decreased ventral striatal activation ^16^ and feedback related negativity (FRN) signals to rewarding stimuli ^27^.

Reward outcome encoding has long been thought as a function of dopamine projections to the frontal-striatal circuit ^28^. However, dopamine replacement does not restore the motivational deficit in the decision to act in apathetic patients ^25^. Furthermore, the results of dopamine replacement treatment strategies have been mixed ^29^ and there is no clear relationship between apathy severity and dopaminergic medication dose ^3^.

These inconsistencies have prompted a re-evaluation of the dopaminergic theory of apathy, particularly in the context of PD. Accordingly, in non-clinical contexts ^30, 31^, many researchers have come to question whether dysfunction within dopamine and its related circuits is critical to the pathophysiology of apathy arising *de novo* in PD^25, 32^. This shift in perspective underscores the complexity of apathy’s neurobiological underpinnings, hinting at the potential involvement of other neural pathways and neurotransmitter systems.

Recent advances in imaging of non-dopaminergic neurotransmitter systems are starting to support this reappraisal, as abnormalities of serotonergic ^32^ and noradrenergic systems ^33^ correlate with the severity of apathy in PD. Serotonin and noradrenaline are known to have overlapping functions with dopamine, including encoding of reward value ^34^ and effort ^35^ . Theoretical and recent empirical evidence, suggest discrete contributions by different projections to the pre-frontal cortex by non-dopaminergic brainstem nuclei in motivated decision behaviour^36, 37^. The activity of these neuromodulators governs whether to weigh up the benefits of either persisting with or changing a course of action to explore and gather new information about the availability of resources. They are also thought to regulate goal directed decisions based upon existing internal models of the world, or to build new ones, based upon new sensory evidence, a failure of which has been implicated in PD-Apathy^38–41^.

Given this context, we tested the hypothesis that apathy in PD is characterised by a decision making signature reflecting a primary failure of outcome monitoring and/ or value-based choice ^19^. To test this, we chose a classical reinforcement learning task, the 4-armed restless bandit ^42, 43^, as its performance relies on the ability to constantly update both the short and longer-term outcomes of each decision. Due to the dynamic and constantly varying payout of each of the “bandits”, performance relies on adaptive behaviour which balances exploitation with exploration ^43–45^. Using a computational model-based fMRI design^46, 47^ we aimed to identify regions of the pre-frontal cortex which underpin apathy in PD.

## Materials and Methods

### Ethics

The study was approved by the local ethics committee (North East Scotland 21/ES/0035). Written consent was obtained from all participants in accordance with the declaration of Helsinki. Seventy-seven participants were recruited.

### Patient group

Fifty-five patients with a clinical diagnosis of idiopathic Parkinson’s Disease were recruited from movement disorders clinics in NHS Tayside, Grampian and Greater Glasgow & Clyde, UK. Diagnosis was confirmed by a consultant neurologist (TG, VM, AM) guided by UK brain bank criteria.

### Control group

Twenty-two -age and sex-matched healthy controls were recruited via the SHARE health informatics register (https://www.registerforshare.org/). Healthy controls were screened for a history of significant neurological or psychiatric conditions.

### Exclusion Criteria

Patients were excluded if they had a diagnosis of Parkinson’s disease Dementia or any other co-morbid neuropsychiatric diagnosis, including Major Depressive disorder. PD patients on anti-depressant therapy in remission from depression were included in the study but could not be under active treatment by a Consultant Psychiatrist. No patient was receiving antipsychotic medication. Two patients were excluded as their Montreal cognitive assessment score was within the abnormal range (MoCA<24).

### Procedure

Participants performed two sessions. The first, “Out of Scanner” session, involved performing the restless bandit task on a laptop, while the second “In Scanner” session had the task performed during fMRI image acquisition. All assessments and tasks were performed with the patients on their usual Parkinson’s medications. If patients had no contraindications for MRI scanning (e.g., metal implantation, claustrophobia, or significant dyskinesia that could lead to image motion artifacts), they underwent two separate sessions on the same day.

### Clinical rating scales

Apathy was assessed using the Lille Apathy Rating Scale (LARS) a questionnaire specifically validated for assessment of Parkinson’s Disease. LARS scores range from -36 to +36 with scores >-22 considered apathetic ^48^. Parkinson’s Disease severity was assessed using part III of the Movement Disorders Society Unified Parkinson Disease Rating scale (UPDRS)^49^ in the ON medication state. Mood and anxiety scores were assessed using the Hospital Anxiety and Depression scale (HADS-A &-D). Cognitive screening was performed using the Montreal Cognitive Assessment (MoCA). Participant demographics are in Table 1. LARS factorial sub-scores in Supplementary Table 1.

**Table 1.** Patient demographics and clinical details. Values expressed as mean ± standard deviation. LARS, Lille Apathy Rating Scale, UPDRS, Unified Parkinson’s Disease Rating Scale, HADS, Hospital Anxiety and Depression Rating Scale, MoCA, Montreal Cognitive Examination. Two-Tailed unpaired T-test significant differences in bold. * Fishers exact test.

|  | Healthy Controls | Parkinson's Disease (PD) | Control versus Parkinson's Disease (p-value) | PD Apathy (LARS>-22) | PD No Apathy (LARS<-22) | PD Apathy versus PD No Apathy (p-value) |
| --- | --- | --- | --- | --- | --- | --- |
| n | 22 | 53 | n/a | 25 | 28 | n/a |
| Age | 62.8±8.5 | 62.8±8.9 | 0.99 | 61.6±11.3 | 63.8±6.4 | 0.39 |
| Gender M:F | 15:7 | 37:14 | 0.78* | 20:3 | 17:11 | 0.06* |
| Apathy (LARS) | -28.1±3.6 | -21.7±8.1 | **<0.001** | -14.0±5.2 | 28.0±2.9 | **<0.001** |
| UPDRS-III | n/a | 30.3±8.7 | n/a | 31.9±8.6 | 28.7±8.7 | 0.25 |
| Levodopa Equivalent dose (mg/24 hours) | n/a | 643±368 | n/a | 608±388 | 673±354 | 0.55 |
| HADS-D | 2.3±2.4 | 5.5±3.7 | **<0.001** | 8.5±2.6 | 3.0±2.4 | **<0.001** |
| HADS-A | 4.9±2.8 | 6.7±4.1 | 0.07 | 8.1±4.2 | 5.5±3.7 | **0.02** |
| MoCA | 27.4±2.0 | 27.6±1.7 | 0.67 | 27.1±2.0 | 28.0±1.24 | 0.07 |

### Experimental design

Participants performed the restless four-armed bandit task ^42, 43^. Subjects were given written instructions on how to perform the task, were told that with each trial they could win between 0 and 100 points and agreed to maximise outcome points.

Each trial started with presentation of four different coloured squares with all four bandit’s levers in the upright position representing the four choice options (Fig.1). Participants made their selection using a four key mini-keyboard (Ecarke-EU) with the colour of each button corresponding to a square presented on the computer monitor corresponding to each of the “bandits”. If a button press was not made within a 1.5 second response deadline, a large red “X” was displayed for 4.2 seconds at the centre of the screen. These trials were designated as missed trials and no outcome feedback was provided. For choices made within the response deadline, the chosen bandit was highlighted with its lever shown depressed and a checker-board pattern appeared at the centre of the bandit’s square. After a 3 second waiting time, this pattern was replaced by the outcome number of points earned on that trial in the centre of the chosen bandit’s square for 1 second. Then the bandit image disappeared and was replaced by a fixation cross until 6 s after the trial onset, followed by a jittered inter-trial interval (Poisson distribution, mean: 2 s (0–5 s)) before the next trial was started. The payout (outcome) schedule of each of the four bandit choices varied according to a decaying Gaussian random walk. We used two instantiations from Daw et. al., (2006) ^42^ for the two experimental sessions and the order of sessions was the same for all subjects.

**Fig. 1.**
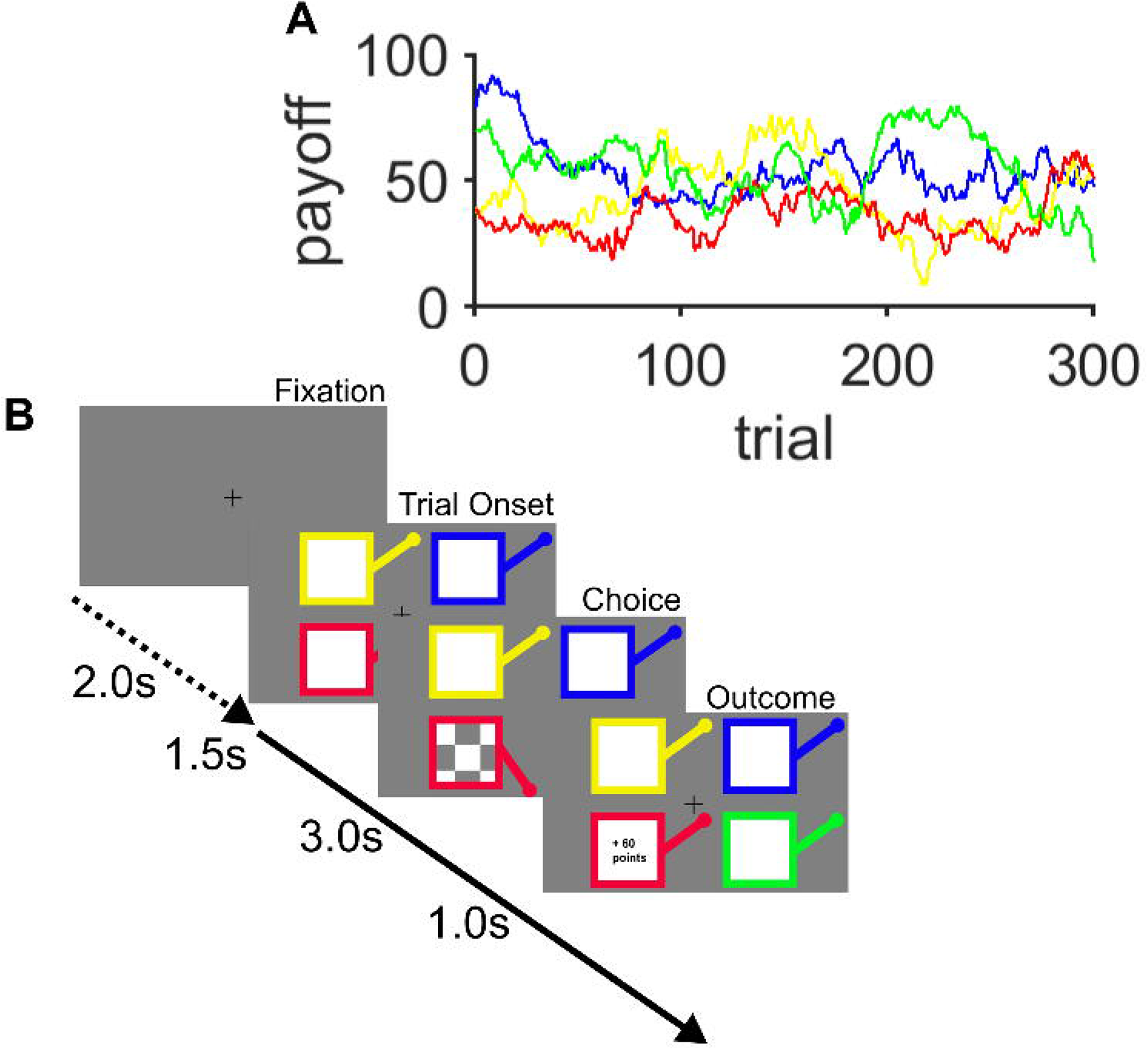
Restless bandit Task. (**A**) Example of the underlying payout (reward) structure across the 300 trials of the task for each of the 4 bandits. The payout varied from one trial to the next by a gaussian walk. (**B**) Each trial has a fixed trial length of 6 seconds with a variable intertrial interval designated by the time between the fixation cross and the onset of the trial (mean 2 seconds). At trial onset, four coloured squares (bandits) were presented. The participant selected one bandit within 1.5 s, which was then highlighted by the bandit lever depressing and a chequer board appearing (Choice screen). After a 3 s delay the, the outcome of the choice, as number of “points” won, was displayed for 1 s. In trials where the participant failed to make a choice within 1.5 seconds the choice screen was replaced by a large red cross (not shown) signifying a missed trial.

During the “In-Scanner” session participants made responses using an MRI-compatible button box. During this session, task images were projected onto a screen visible to the patient inside the MRI scanner. The task was implemented using the MATLAB (R2021; MathWorks, Natick, MA) running Psychophysics Toolbox (version 3.0.12) ^50^.

Of the 53 patients who participated in the out of scanner session, 40 (20 with apathy, 20 without) also agreed to an In-Scanner session, along with 20 of the healthy controls. One patient in the apathy group was unable to perform the task in the scanner and was exclude from final analysis.

The task consisted of 300 trials. Participants were given short breaks after every 75 trials to improve concentration and task engagement.

### Analysis of behavioural performance

Model free metrics of behavioural performance in the task included best bandit choice probability, decision time and the probability of missing a trial. Decision time was defined as the time between the four bandits being presented and the participants’ button press on that trial. Where a measure of behavioural performance is expressed as a probability, this was achieved by dividing by the total number of trials correctly executed within response deadline, either in a 50-trial block, or by dividing this by the total number of responses made across the whole task.

### Statistical analysis

We used a mixed-design ANOVA with a fixed effect (between subject) of group with three levels, (PD-apathy, PD-no apathy, healthy control and a random effect (within subject) variable of within task block with (six, 50-trial blocks). All results are reported as mean values ± S.E.M.

### Computational modelling of decision making

To understand the decision-making process further in PD-apathy we fitted 8 computational model variations of the decision process.

Each model variant used one of two learning rules, (Delta rule, or Bayesian learner) combined with one of four choice rules: i) Softmax (SM), ii) Softmax with exploration bonus (SME), iii) Softmax with perseveration bonus (SMP), (iv) Softmax with exploration and perseveration bonuses (SMEP). Posterior parameter distributions were estimated for each subject for each of the free parameters specific to the learning (a in the Delta learning rule) and choice rules for each model variant (/3, cp, p). Details of the model and fitting procedure are provided in Supplementary Material.

### fMRI methods

Details of fMRI sequence acquisition are in Supplementary Materials. The fMRI analysis was performed on three distinct participant groups: healthy controls, patients diagnosed with Parkinson’s disease who exhibited symptoms of apathy (PD-apathy), and patients with Parkinson’s disease without apathy symptoms (PD-no apathy). We conducted a first-level analysis (detailed in Supplementary Materials) by creating a general linear model (GLM) for each participant. This was done separately for each group across the four blocks of the In-Scanner task.

Utilizing a second level random effects approach, the subject- and group-specific contrast images for each first-level regressor were submitted to a full factorial model in SPM12. For each contrast specific second level analysis, a T-contrast image was generated and tested for the main effect of that contrast over all subjects for each of the groups.

We performed whole-brain analyses of both activation within groups and between group contrasts and report activations and between group contrasts surviving cluster-level FWE correction at *P* < 0.05 (indicated with p_cluster FWE WB_), corresponding to a simultaneous requirement for a voxel threshold of *P* < 0.001 and a minimum cluster size of 10 voxels. Region of interest analyses were performed using a 10mm sphere centred on a peak voxel of interest (indicated with p_peak FWE SVC_).

## Results

### Patients with apathy are less likely to choose the best bandit

We acquired choice behaviour from 53 Parkinson’s patients (25 with apathy, ”PD-apathy”, and 28 without, “PD-no apathy” 22 age- and sex-matched healthy controls, “HC”, performing a four-armed “restless“ bandit task (Fig. 1). Patients with and without apathy had comparable levels of motor disability and medication status (Table 1).

In the Out-of-Scanner session, patients with PD-apathy learned to choose the best of the four bandits above chance levels (P(best bandit):0.54±0.04) but were less likely to choose the best compared to the PD no-apathy (P(best bandit):0.64±0.03) or HC (P(best bandit):0.65±0.03) groups; Main effect of Group F(2,344) = 5.54, p=0.005, (Fig. 2A). The probability of choosing the best bandit correlated with each patient’s apathy severity (Fig. 2C) as indexed by their total LARS score (r = -0.43, n = 56, p = 0.001).

**Fig. 2.**
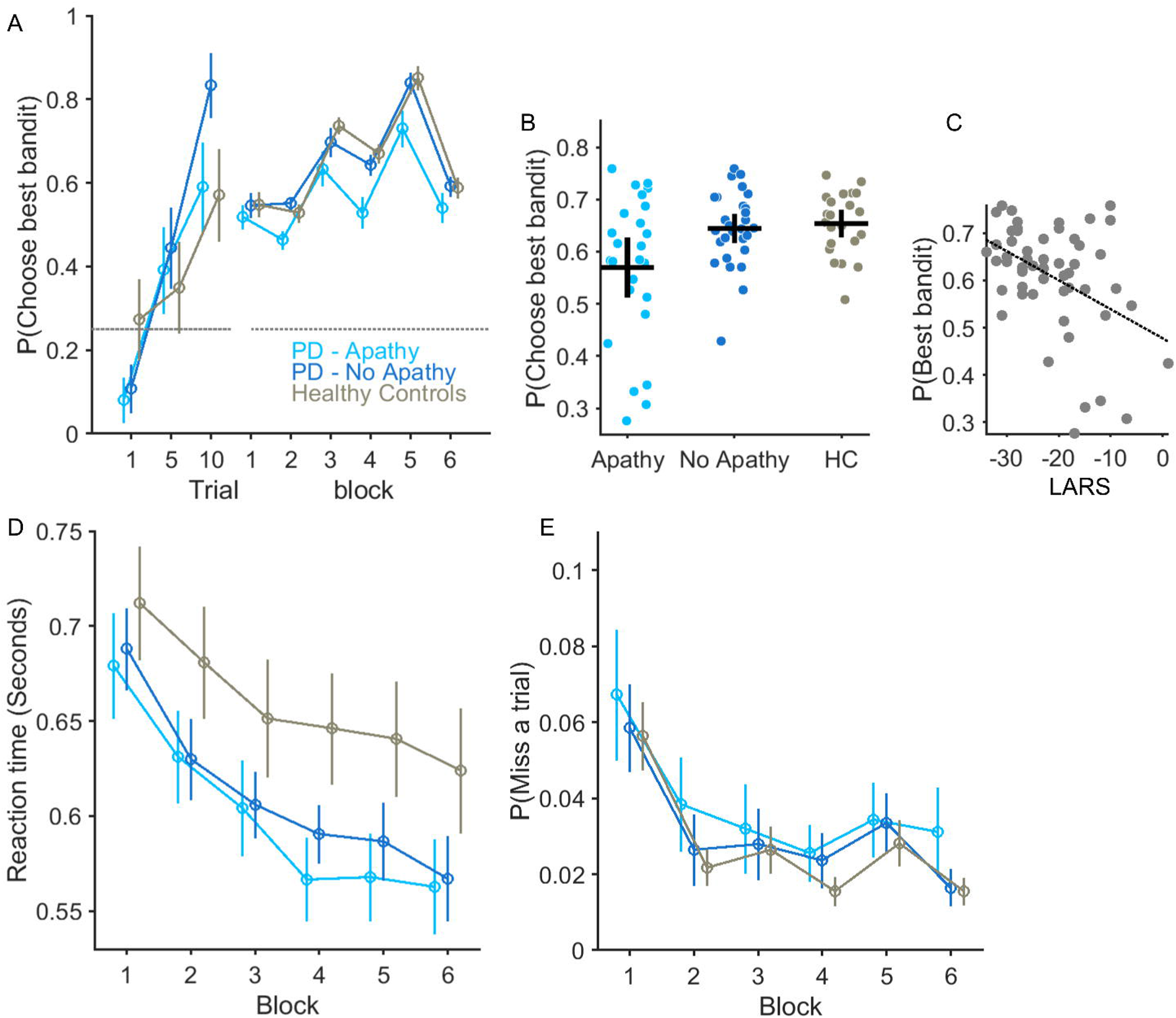
Bandit performance and relationship with Apathy Severity. The average probability of choosing the bandit with the highest payout, P(Choose best bandit) is plotted in the three groups of participants. (**A**) The increase in average values of best bandit choice plotted for in trials 1, 5 and 10 confirm learning in all three groups from an initial random choice to above chance levels (horizontal dashed line). Best bandit choice performance across six, 50-trial blocks of the task was reduced in the PD-Apathy group. Vertical lines S.E.M. In **(B**) each circle represents average best bandit choice probability across the task for an individual subject with the horizontal and vertical bar represents the group mean and 95% confidence limits. Apathy severity, measured by increasing LARS score (more positive values represent higher levels of apathy) correlated with individuals’ ability to choice the best bandit (r = -0.43, p = 0.001). Each group’s average reaction time over the 6 task bins in (**D**) and likelihood of not making a response (**E**) (missed trial) was not affected by apathy.

Impaired best bandit choice in the task could not be clearly explained by diminished engagement in the task, as the decrease in decision time seen across blocks in both HC and PD no-apathy groups was also observed in the PD-apathy group (Main effect of block F(5,344) = 26, p<0.001, block*group interaction F(10,344) = 0.48, p=0.75 Fig. 2D). There was no significant difference in average decision times between groups over the course of the task (Decision time: PD-apathy, 0.60±0.02 seconds, PD-no apathy 0.61± 0.02 and HC 0.65±0.03) or effect of Group on Decision time (F(2,344) = 1.8921 p = 0.15). Moreover, the apathetic patients did not miss significantly more trials than patients without apathy or healthy controls (Fig. 2E). (Probability of missing a trial: PD-apathy 0.03± 0.01 PD-no apathy, 0.03±0.01, HC 0.02±0.006, Effect of Group F(2,344) = 0.5, p=0.6)). We reproduced the same behaviour in the In-Scanner session (Supplementary Material & Supplementary Fig.1).

### Explore-exploit trade off predicts individual apathy severity

This analysis confirmed that PD-apathy was correlated with the ability to monitor the outcome of a fluctuating payout requiring identification of the most rewarding choice. We further hypothesised this could arise from three mutually exclusive mechanisms.

First, patients with apathy might use a *perseverative* strategy that minimises the cognitive effort, by making choices irrespective of the perceived value of an option ^51^. Second, apathetic patients may employ an overtly greedy choice strategy. Whilst this may initially seem advantageous, it reduces the information gained from non-greedy, or exploratory, choices resulting in poorer decision flexibility ^52, 53^. Finally, if the neural representation of decision value is degraded (or encoded but the information disregarded), behaviour should be characterised by choice policy of heightened *exploration* which reflects heighted uncertainty, about which of the options is best ^54^.

To further investigate these mechanisms, we utilized a model of the neural decision-making process. Our goal was to approximate how the brain encodes the expected value of a decision. Consistent with the winning model (Bayes-SMEP, Supplementary Fig.2 A) capturing decision making in the task robustly, model parameters could still be recovered (Supplementary Fig.2 B-D) from synthetic choice data generated from simulated choices. These also overlapped with the experimental choices from each group (Supplementary Fig. 3).

By modelling the value of each of the four options throughout the task, we were able to categorise each choice into three categories: exploitative, directed exploratory, and random exploratory choices.

An exploitative choice refers to the selection of the option perceived to have the highest expected value. A directed exploratory choice is one made towards one of the three less valued options, particularly the one not selected recently^53^. Random exploratory choices, on the other hand, involve the selection of a less valued option regardless of current knowledge or uncertainty about their payout^53, 55^.

Patients with apathy made a significantly higher proportion of random exploratory choices (probability of random exploration, or P(RE)), (Fig. 3 D-F). The P(RE) was in the PD-apathy group = 0.23±0.02, PD-no apathy= 0.15±0.010, and 0.13±0.009 in the HC(Main effect of Group F(2,344) = 8.69, p<0.001). PD-apathy patients made fewer exploitative choices through the task than non-apathetic PD patients (Fig.3 A&B) and HC (P(Exploit): PD-apathy = 0.62±0.03, PD-no apathy = 0.72±0.015, HC = 0.73±0.018, Main effect of Group F(2,344) = 6.31, p=0.002). The proportion of directed exploratory choices did not differ between groups (P(DE); PD-apathy = 0.13±0.018, PD-no apathy = 0.11±0.009, HC = 0.13±0.01, Main effect of Group F(2,344) = 0.86, p= 0.42).

**Fig. 3.**
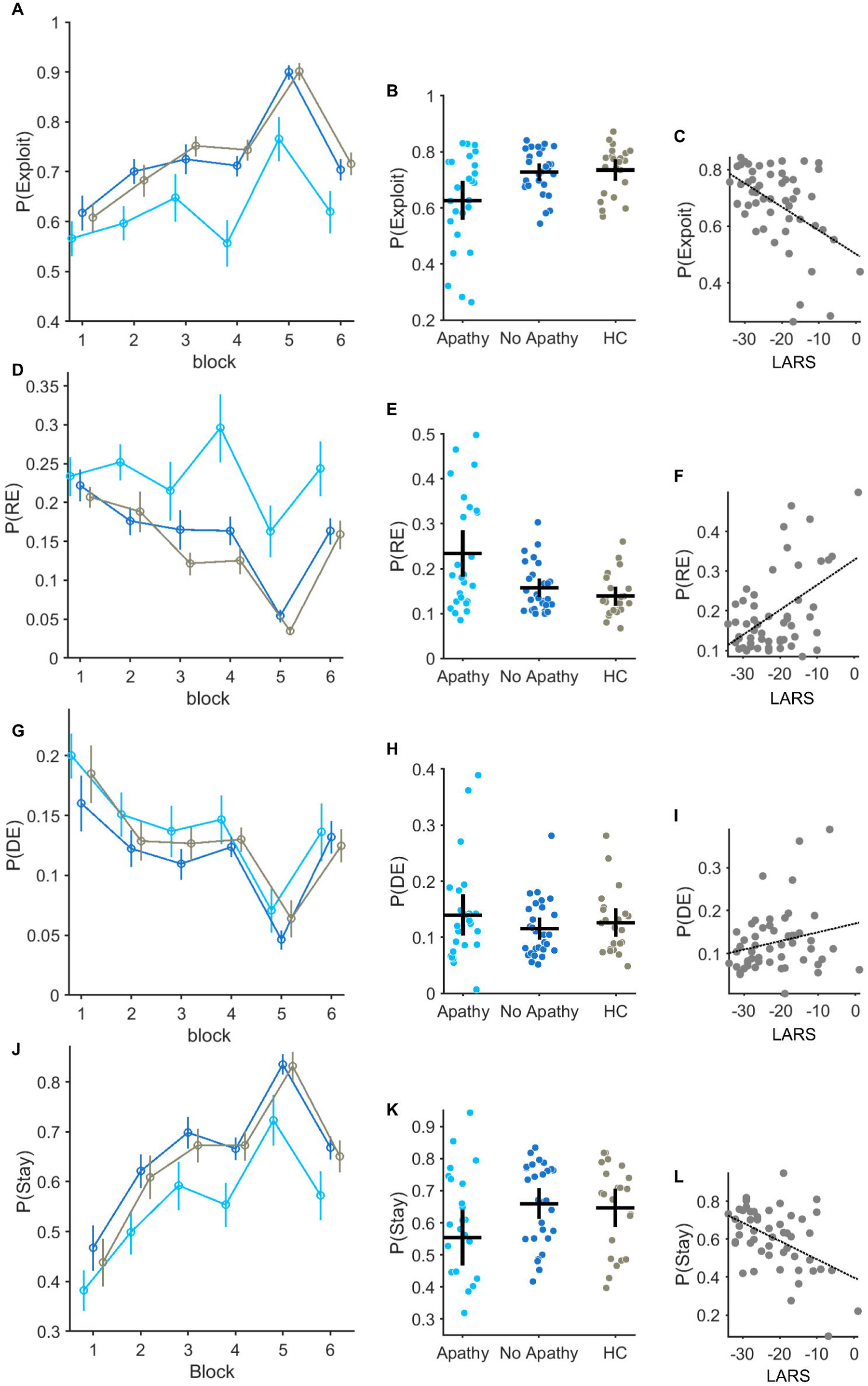
Decision types between groups across the task and relationship with Apathy severity. **(A-C)** Probability of making an exploit P(Exploit) choice plotted across six-50 trial bins for each group tested (**A**). Error bars represent S.E.M. **(B)** individual P(Exploit) across the whole task is represented by each circle. Group average and S.E.M is illustrated by the vertical and horizontal bars. Correlation between P(Exploit) and Apathy severity (increasing LARS Score) in (**C**) r = -0.50, p<0.001. The same analysis applied to the probability of making a random exploratory P(Explore) choice **(D-E)** and relationship with individual apathy severity (**F**) r = 0.47, p<0.001. P(DE) is the probability of making a directed exploratory choice (**G-I**) and P(Stay) the same choice on two consecutive trials (**J-K**). (**L**) Correlation between P(Stay) and Apathy severity r = -0.47, p<0.001.

The proportion of both exploratory and exploitative choices correlated with the severity of apathy in individual patients, with P(RE) versus LARS recording rho(52) = 0.47, p<0.001, and P(Exploit) versus LARS recording rho(52) = -0.50, p<0.001 (Fig. 3C&F).The proportion of perseverative choices, a model-free index of the likelihood of making the same choice on at least two trials, was also lower in the PD-apathy group, P(Stay) PD-apathy = 0.55±0.04, PD-no apathy = 0.65±0.036, HC = 0.64±0.04, Main effect of Group F(2,344) = 3.22, p=0.04), (Fig 3 J&K) and correlated with the LARS score, rho(52) =-0.47 p<0.001 (Fig. 3L).

This different decision signature between the patient groups was also reflected in the model parameter estimates. The inverse temperature parameter, /3, proportionately scales the extent a choices value estimate is used in the decision. Consistent with reduced exploitation in PD-Apathy group, /3 values were significantly lower: posterior difference in mean, [Contrast: PD-Apathy minus PD-no apathy], M_diff_ = -0.02 [-0.028, -0.015]. The exploration cp, and perseveration bonuses, p, which govern the proportion of directed exploration and perseverative choices were also lower than the PD-no apathy group: cp, M_diff_ = -0.40[-0.69, -0.12], p,M_diff_ = -3.57 [-4.88, -2.27] (Supplementary Fig. 4 4B,E &H, Supplementary Table 2). Apathy severity correlated with the individual subjects parameter estimates, /3, rho(52) = -0.41, p=0.002, p, rho(52) = -0.41, p=0.002 but not the exploration bonus cp, (Supplementary Fig. 4 C&I).

### fMRI signal of outcome encoding is blunted in apathy

Our analysis of the brain imaging data was motivated by two explanations for our apathetic patient’s behaviour. Could a failure to monitor the outcome of their actions, reflected in a shift from exploitation to exploration, be driven by a pure disorder of encoding the outcome of their decisions in the task? If the precision of the outcome’s value signal is degraded ^40^, exploration is likely to be a *passive*, secondary consequence of greater decision noise ^52^. Alternatively, an inability to exploit knowledge gained from learning the value of each option could arise from a failure of using this at the point at which the decision is being made.

We proceeded to analyse the brain activity at the point in each trial at which the outcome was received and looked at the fMRI signal correlation with this payout on each trial. Replicating previous studies in healthy controls ^42, 43, 56, 57^, we identified activity in the ventromedial prefrontal cortex (vmPFC) in our age matched HC group (left vmPFC: peak voxel: *x*, *y*, *z* = -6, 34, −8, *T* = 5.25, p_cluster FWE WB_ < 0.001). However, neither PD patient groups (PD-apathy and PD-no apathy), demonstrated clusters surviving whole brain correction (Supplementary Table 5, Fig. 4).

**Fig. 4.**
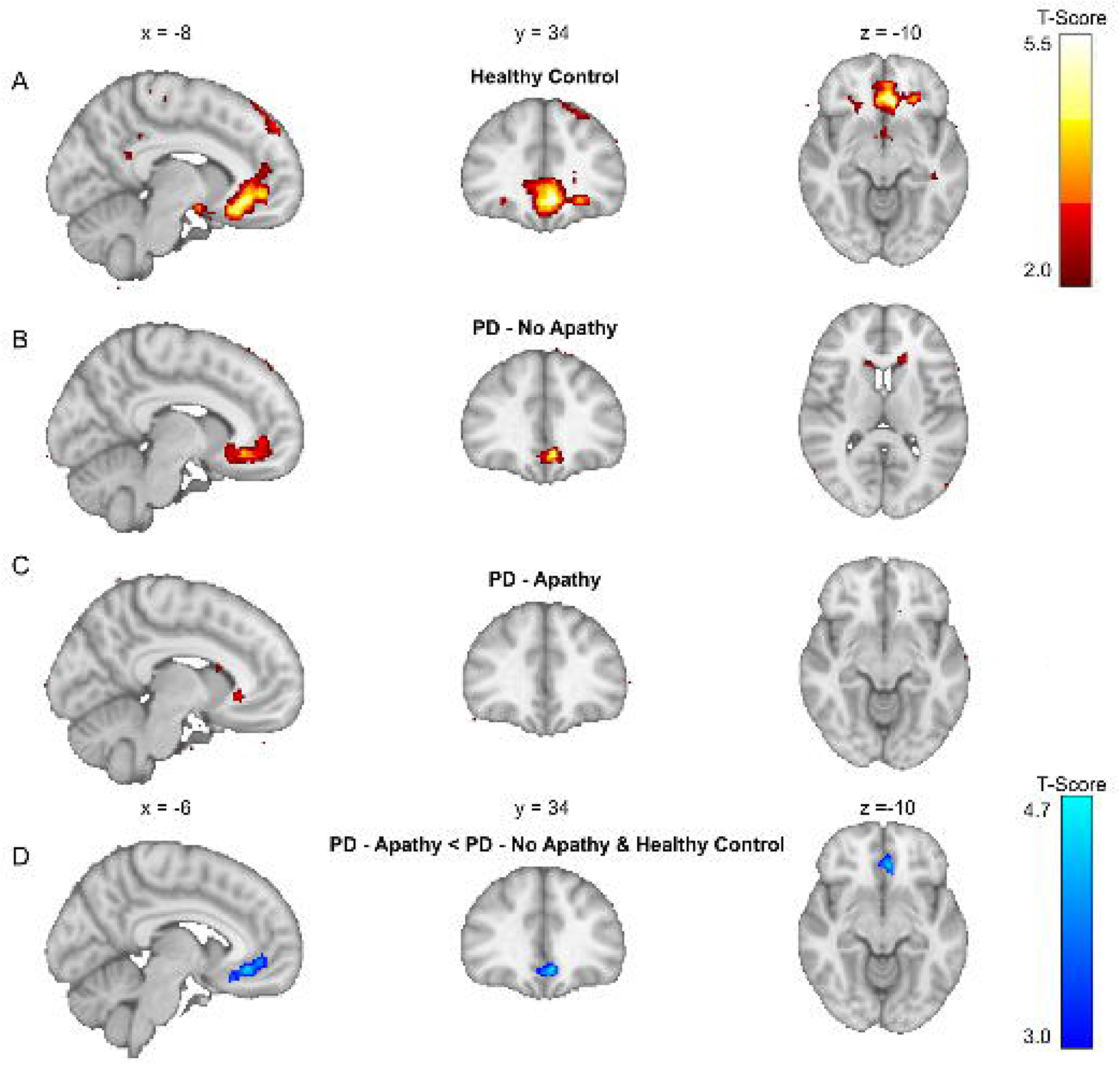
fMRI BOLD correlates of the outcome signal in PD-Apathy and Non-Apathetic groups. **(A)** Peak activations in healthy control (HC) group at the point of feedback of the choice outcome (payoff) time demonstrated a significant cluster in left ventromedial prefrontal cortex (vmPFC) after FWE whole brain correction at p<0.05 (*T*□=□5.25, pcluster FWE WB□<□0.001). **(B**) Activations in PD – No Apathy group did not survive correction at a whole brain level but were present in vmPFC when analysed with small volume correction using a region of interest analysis centred on the peak activation in the HC group (*T*□=□4.51, p_peak FWE SVC_□=□0.004). No significant clusters activity survived whole brain or small volume correction in the PD-Apathy group (**C**). Contrast analysis between groups did not demonstrate any difference in outcome signal activations between the PD-Apathy and PD-NoApathy groups (result not illustrated). However, combining the PD-No Apathy and HC groups confirmed a significant reduction in the outcome signal in left vmPFC in the PD-Apathy patients (**D**) (T = 4.60, p_cluster FWE WB_□= 0.01).

Using a region of interest (ROI) analysis with small volume correction, using the peak vmPFC activations from our HC group, we confirmed vmPFC activation in the non-Apathetic PD group (left vmPFC peak voxel: *x*, *y*, *z* = -6, 34, -10, *T* = 4.51, p_peak FWE SVC_ = 0.004) which was not present using the same approach in the PD-apathy group. However, contrast analysis (PD-apathy<PD-no apathy) using the same ROI, did not confirm a significant reduction in the outcome signal between groups. Combining the PD-No Apathy and HC group in a contrast analysis with the PD-apathy (PD-apathy< PD-no apathy & HC) did reveal a significant reduction in outcome signal including within left vmPFC: peak voxel: x,y,z = -6,34,-10, T = 4.60, p_cluster FWE WB_ = 0.01), (Fig. 4D) . We also performed regression analysis of the individual LARS score in both PD groups with the payout signal at the outcome time. No significant clusters survived whole brain correction. Therefore, despite a reduction in this signal in PD-apathy patients, a difference in the encoding of the outcome signal could provide a singular explanation for their level of apathy, or equally a shift away from exploitation to exploration in the apathetic group.

### Individual apathy severity correlates with cortico-thalamic activation at the decision time

Next, we considered brain activity at the decision-making events, focusing on distinguishing activations between exploratory and exploitative choices at trial onset (Supplementary Fig. 5, Supplementary Fig.6, Supplementary Table 4 for in-scanner model fitting). Our findings were largely consistent with previous research, with the brain demonstrating distinct activity patterns for each choice type. In the healthy control (HC) group, exploratory trials were characterised by recruitment of occipito-parietal regions, including the calcarine cortex (CC), intraparietal sulcus (IPS), and superior parietal lobules (SPL) (peak voxel right CC: x,y,z = 16,-88, 10, T = 7.57, p_cluster FWE WB_ < 0.001). This activity was in conjunction with activations observed in areas previously linked with exploration ^42, 43, 56, 57^ : the bilateral thalamus/midbrain (TH), anterior insula (AI), middle frontal gyrus (MFG), and supplementary motor area (preSMA/dorsal anterior cingulate gyrus(dACC)). Peak voxels in right TH : x,y,z = 8,-24,-4, *T* = 4.61, p_cluster FWE WB_ < 0.001, right AI : x,y,z = 30,24, -4, *T* = 5.50, p_cluster FWE WB_ < 0.001, right MFG : x,y,z = -42, 8, 32, *T* = 5.55, p_cluster FWE WB_ < 0.001, Left preSMA/dACC : x,y,z = -8,16, 46, *T*=5.06, p_cluster FWE WB_ < 0.001 (Fig. 5A & Supplementary Table 6).

**Fig. 5.**
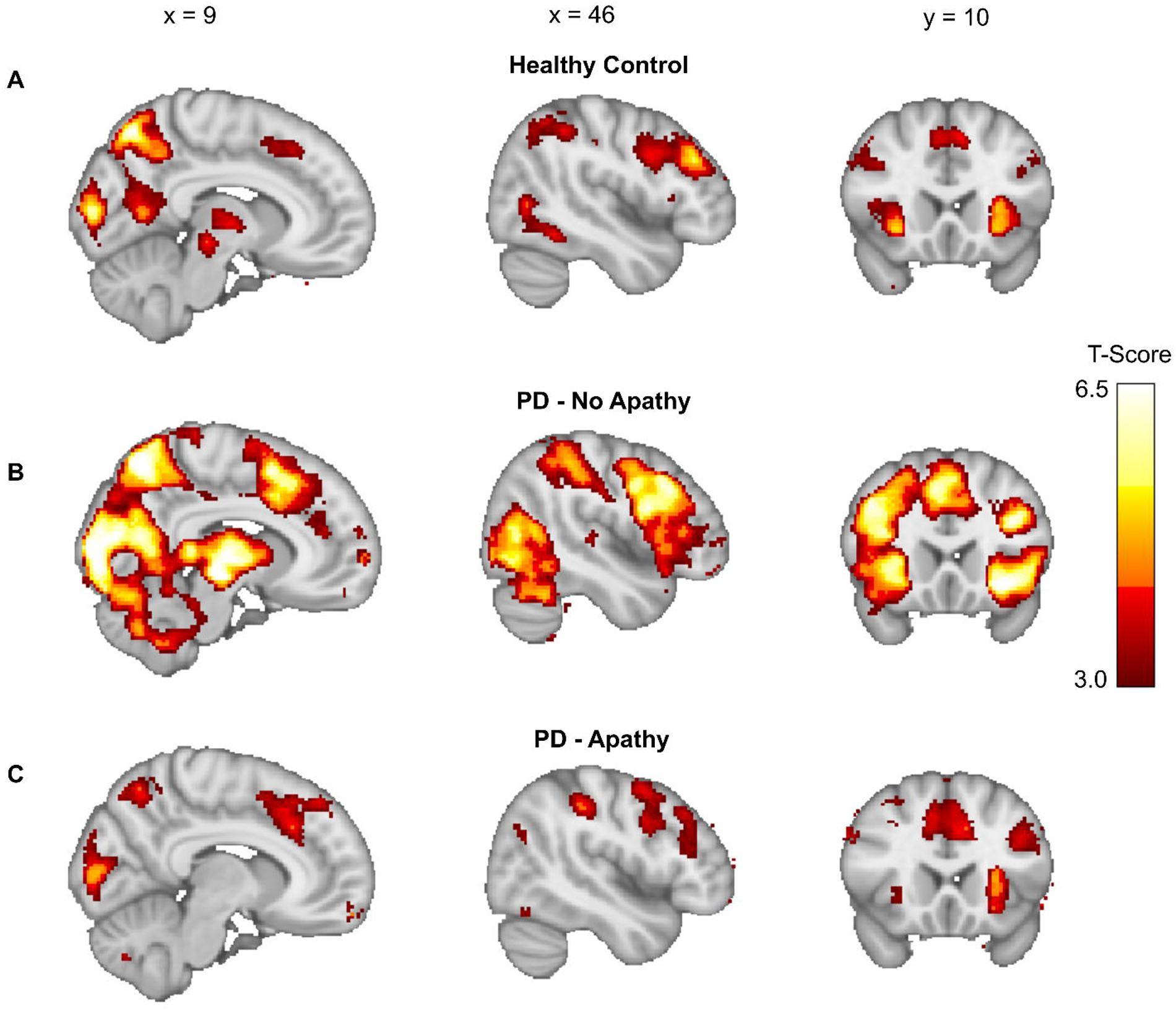
fMRI BOLD signal correlates at the decision time during exploratory choices. **(A)** Peak activations in healthy control (HC) group at the point of decision making when an exploratory choice was made activated parieto-occipital, thalamic and both medial (preSMA/dACC), lateral pre-frontal (DLPFC) and bilateral anterior insula regions. Corresponding activation in PD-No Apathy (**B**) and PD-Apathy **(C**) groups. Significant clusters were defined as those surviving FWE whole brain correction at p<0.05 (See Supplementary Table 4).

In contrast, exploit decisions in the HC group, activated more limited regions consisting of superior frontal gyrus (SFG), and vmPFC/ subgenual cingulate. Peak voxels left SFG: x,y,z = -18,52,22, T = 4.63 p_cluster FWE WB_ < 0.003; left vmPFC: x,y,z = -18,36,4, T = 4.13 p_cluster FWE WB_ < 0.001, (Supplementary Fig. 6). No significant clusters survived whole brain correction during exploit trials in either the PD-apathy or PD-no apathy groups (Supplementary Table 6) and there was no significant difference in the activations between these groups.

The same clusters of activity during explore choices were noted in both the PD-apathy (Fig. 5C) and PD-no apathy (Fig. 5B) groups within MFG, dACC, Left FP, and posterior occipito-parietal regions including CC, IPS, SPL and FP (Supplementary Table 7). Decision to explore activations were generally more marked in PD-no apathy group than in both the HC and PD-apathy groups.

Comparing activations between groups using a second level contrast (PD-no apathy >HC) confirmed additional recruitment of pre/post central gyrus (PCG), Cerebellum (CB), Inferior Frontal Gyrus (IFG) and Frontal Pole (FP). Peak voxels PCG: x,y,z = 0,-30,68, *T* = 5.52, p_cluster FWE WB_ < 0.001, left CB: x,y,z = -26,-52,-42, *T* = 4.28, p_cluster FWE WB_ < 0.001, IFG: x,y,z = -32,-26,14, *T* = 4.35, p_clusterFWE WB_ < 0.001 and FP but not in TH, MFG, AI or preSMA/dACC (supplementary Table 8).

Further contrast analysis between the PD-apathy and HC groups (PD-apathy< HC) signal was comparable levels to those of HC as no significant clusters were above threshold following whole brain cluster level correction (Supplementary Table 7).

In the PD-apathy group, exploratory choices were associated with intact activity across the same bilateral prefrontal (MFG), preSMA/dACC and anterior insula regions as non-apathetic participants. However, there was no comparable activation of the thalamic/midbrain and occipitoparietal regions seen during exploratory choices in the HC and non-apathetic PD groups.

This absence of thalamic/midbrain and occipitoparietal activation in the PD-apathy group was verified by a contrast analysis comparing the PD-no apathy and PD-Apathy groups (PD apathy< PD no apathy). Post contrast analysis, clusters that survived whole-brain FWE correction were identified in the bilateral thalamus, bilateral pre-post central gyrus, intraparietal sulcus, bilateral cerebellum, and calcarine cortex. Peak voxels right TH : x,y,z = 14,12,2, *T* = 5.37, p_cluster FWE WB_ < 0.001, right PCG : x,y,z = 6,-32,-72, *T* = 4.67, p_cluster FWE WB_ < 0.001, left IPS : x,y,z = 26,-66,-28, *T* = 4.46, p_cluster FWE WB_ < 0.001, left CB : x,y,z = -10,-58,-44, *T* = 4.57, p_cluster FWE WB_ < 0.001 right CC : x,y,z = 26,-52,-4, *T* = 4.89, p_cluster FWE WB_<0.001 (Fig. 6 & Supplementary Table 8).

The activity at the decision time during explore choices was also found to correlate with apathy severity in the same right thalamic cluster identified in the between group contrast analysis (k = 27, peak voxel right TH: x,y,z = 14,-12,2, T = 5.44 p_cluster FWE WB_ = 0.005. Fig.6D).

**Fig. 6.**
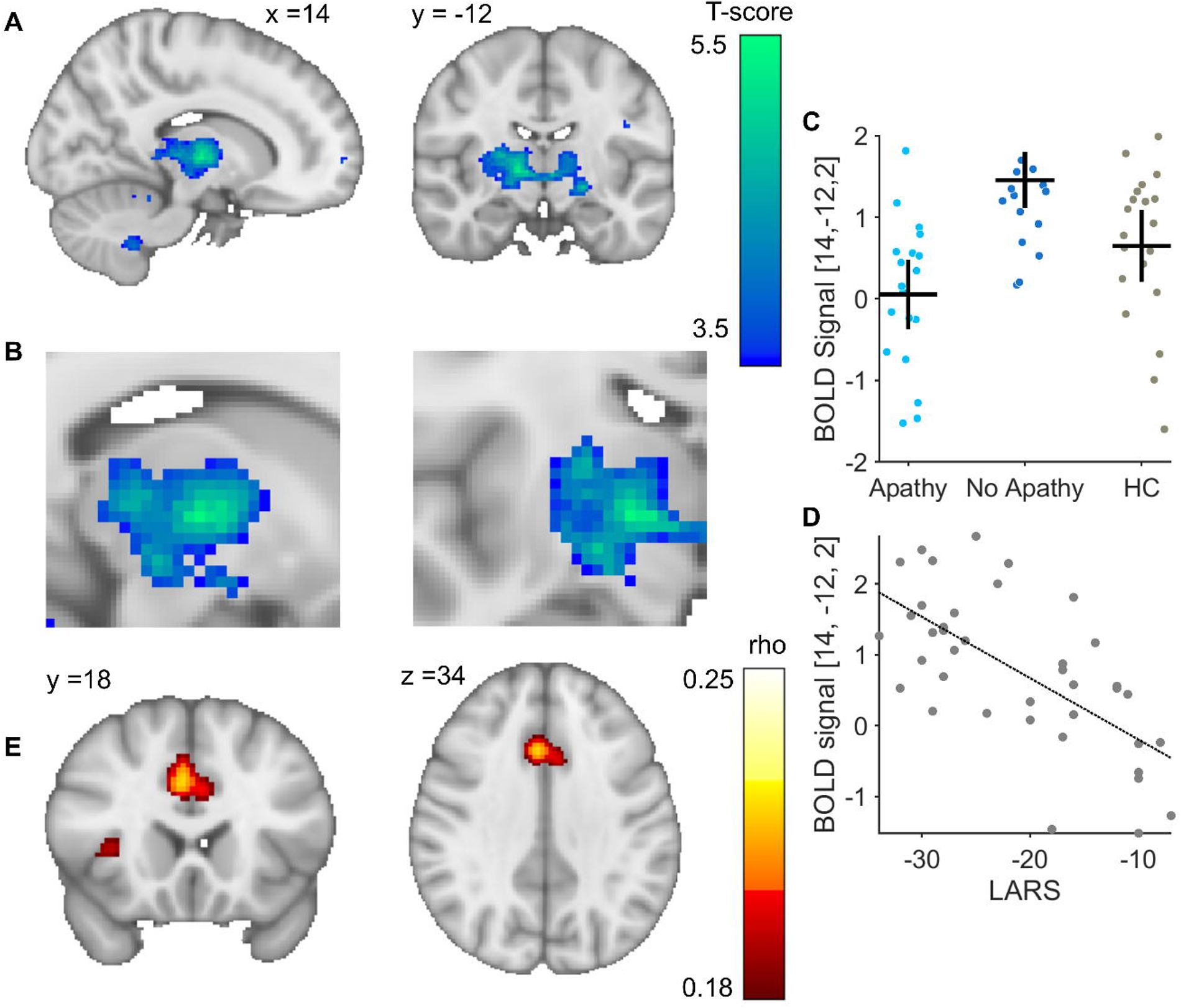
fMRI BOLD contrast differences the decision time during exploratory choices. **(A &B)** Peak contrast difference in activation between the PD-Apathy and No-Apathy within the right thalamus at the point of decision making when an exploratory choice was made. BOLD signal activations at this peak voxel plotted for each subject and group (**C**) and correlation with individual Apathy severity (**D**). Functional connectivity analysis at this voxel using normative functional resting state connectivity showed peak correlation between the thalamus and dACC/ bilateral AI (**E**).

Finally, to identify the cortical functional connection between this thalamic voxel that was a predictor of the patient’s apathy severity, we used *neurosynth* [4, 20,33]. This confirmed peak functional connectivity at x,y,x = 3,18,36, rho = 0.22 corresponding to dACC and bilateral AI, x,y,x =35,7,-1 & – 34,5,-1, rho = 0.23 (FDR corrected for multiple comparisons p< 0.05, Fig. 6E).

## Discussion

We assessed the mechanisms of demotivated behaviour characterised as apathy in Parkinson’s disease, using computational modelling of decision-making behaviour combined with event-related functional MRI. The core results of this study are three-fold. First, patients with PD-apathy are less able to monitor and chose the best option when faced with outcome uncertainty. Second, they use an exploratory decision strategy which is best explained by impairment of incorporation of the neural representation of an options value into decision-making. Third, a loss of compensatory neural circuits implicated in decision evaluation, predicts whether apathy is manifest in PD or not.

Reward insensitivity as a core feature of PD-apathy has been observed in several previous studies ^16, 26, 27^.Our results extend this, whilst simultaneously addressing the question of where in the neuronal processes that leads to motivated (and de-motivated) decisions, reward sensitivity is lost. We found behavioural evidence of reward insensitivity in apathetic patients in the restless bandit task. The ability to choose the most rewarding option was a strong predictor of their apathic status. As revealed from computational modelling, PD patients with apathy make decisions that are indifferent to the learnt expected value of the different options. In modelling terms, this was reflected by the reduction in the reward sensitivity parameter, /3. By multiplying the value estimate for each option learned from one trial to the next, low levels of /3 drive up the proportion of non-greedy, “exploratory” choices to the lower value options. In isolation, this analysis does not explain where the insensitivity to reward occurs, and by extension, its degradation, or the disregard of this learnt neural value representation. We were motivated to study learning in this task because it requires high levels of cognitive demand in both monitoring and learning from the outcomes of actions. At a neural level, to track and identify best option, relies on a reliable and constantly updated representation of the outcomes value (and presumably the relative longer running reward average to compare this too).

One explanation for our results, is therefore, a failure of *encoding* the value of the chosen option within the brain ^58, 59^. In non-human primates, neural firing rates correlate with the outcome received within the vmPFC ^60^. We reproduced the fMRI correlate of this outcome encoding described previously in our age-matched HC group in these regions. While this signal was blunted in both PD groups, it wasn’t a clear discriminator of their apathetic status. Caution is required in interpreting this signal, as this was less robust in the HC group than that previously reported. This may be a reflection of the smaller sample sizes in our groups, but is equally likely to reflect a disease- and age-related pre-frontal dopaminergic denervation ^61, 62^. Combining both the PD no-apathy with the HC group we were able to demonstrate significant blunting of this signal in the PD-apathy patients. Therefore, impairment in the encoding of a choices outcome is a feature of PD-apathy which is additional to that which occurs as a consequence of Parkinson’s disease^63^. However, as impairment of this signal did not correlate with an individual’s apathy severity, we would argue that this is unlikely, in isolation, to be the primary explanation for their demotivation.

At the decision time, we reproduced previously described fMRI correlate of exploration indexed by cortical activations including FP, IPS, AI, preSMA/dACC in all three groups. Exploitative choices activated vmPFC/subgenual cingulate regions in the age-matched control group only. These regions have been implicated in the encoding of the expected value of chosen options ^64–66^. Despite no equivalent activation in both patient groups, paralleling the disease-related blunting of the outcome signal, we were not able to demonstrate any between group differences. In studies of younger HC, exploitation activates a more extensive set of regions (lateral OFC, Posterior cingulate, hippocampus and precuneus ^42, 43, 56, 57^ which overlap with those of the default Mode Network (DMN)^67^. Age- and disease related decline in the DMN signal in prefrontal cortex may explain the absence of a significant between group difference as a consequence of reduced signal – to noise ^68, 69^.

The overlap between activations during exploitation with regions in the DMN have led to the view of exploitation as a decision choice with lower cognitive and attentional demands ^70^. Exploration activates regions whose functions are synonymous with cognitive control. For example, the AI/dACC form nodes of the salience network (SN), which heightens the detection of behaviourally relevant stimuli, response selection and conflict monitoring^71^. The IPS is thought to serve as interface between prefrontal regions and motor output, initiating the behaviour necessary to explore alternative actions ^42, 72^. Finally, the FP tracts the uncertainty of unsampled options to trigger switches from exploitative to exploratory behaviour ^57, 73^. Recruiting these regions may explain why learning (and learning rates) increase during exploration ^39, 74^. Why then, do our apathetic patients appear to perform worse in our task? We would argue that despite classification of these choices exploratory, they are not analogous to exploration that occurs in health and do not reflect heighted cognitive control. Firstly, exploration is costly ^21, 75^. Intuitively, it seems unlikely that patients with increased sensitivity to effort, would overly rely upon a decision strategy which is comes with additional cognitive cost. This is also at odds with their clinically demotivated state, one manifested by indifference to novelty seeking and a withdrawal from salient external events within their wider environment ^11, 76, 77^.

Second, we found no difference in cortical activations during exploration in the PD-apathy and HC groups or additional activity in PD-apathy that was not present in the non-apathetic PD patients. This would argue against exploration being actively driven by a gain of neural function in which one or more brain regions is actively promoting this behaviour. This, for example, might be driven by uncertainty about the environment ^47^, or greater sensitivity to opportunity cost^78^.

We would argue that rather than being an active process to either seek out or learn new information, the shift from exploitation to exploration in PD–apathy, is a *pathological* signature of increased decision noise. Indeterminate, random selection rules are efficient strategies for exploration and are necessary for optimal adaptive choice ^53, 79^. Sub-optimal, “non-greedy”, choices are a feature of normal decision making in health and are thought to arise from limitations in the brain encoding of a decision’s value ^52, 54, 80^. By degrading the precision of outcome encoding or the encoding of an action’s expected value, pre-frontal dopaminergic denervation could explain increased decision noise in PD^81, 82^. This loss of outcome and/or value encoding occurring independently of whether or not the patient has apathy^63^.

We interpret the finding of the increased activation of the explore circuit in PD-no apathy group as evidence for a compensatory mechanism that may protect against manifestation of apathy. In support of this, was the finding that the peak contrast difference between the PD groups correlated with individual apathy severity (LARS score). A compensatory mechanism is also more likely given that there was no difference between the PD-apathy and HC group activations, but increased activity in these regions was observed in the PD-no apathy group relative to the healthy controls. Connectivity between the medial and anterior thalamic nuclei and dACC positively correlates with enhanced cognitive performance and goal directed behaviour in uncertainty ^83, 84^. Additional recruitment of this circuit could preserve exploitation in our PD no-apathy group as this included cortical areas of the SN and thalamus with peak functional connectivity with the dACC.

Alternatively, the same over-recruitment of these regions during exploration, can also be viewed as an equivalent *deactivation* in the same regions during exploitation. During exploration, neural firing in pre-frontal cortex adopts a state transition into an indeterminate, disorganised and non-coding state^85^. It may become imperative, when the brain loses the ability to encode value or outcome with precision, to suppress activity in circuits that may naturally augment additional neural decision noise. By over-suppressing activity in these (*explore*) regions during *exploit* decisions, this would limit their influence to promote decision noise being expressed into behaviour. This would support the possibility that In Parkinson’s apathy, the combination of *both* a loss of reward encoding and loss of mechanisms which override decision noise leads to demotivated behaviour. In our apathetic patients, the behavioural expression of this combined loss of function, is manifest in decisions to explore rather than exploit.

### Study Limitations

Could a simpler explanation of demotivated task engagement explain the behaviour of our apathy patients? For example, might heightened exploration be explained by an apathy related indifference to task performance to allocating cognitive costs. Against this is that we found no clear difference in either the decision time or proportion of missed trials in the PD-apathy and no apathy groups. Furthermore, at the decision time, explore choices in the Apathy group activated prefrontal cortical regions including dACC and DLPFC, regions whose activation indexes engagement with task complexity ^86^and working memory ^87^.Their activation to levels in comparable to that seen in the HC group would make a deficiency in the allocation of cognitive control an unlikely explanation for their poorer task performance. Demotivation leading to task disengagement may also arise through heightened sensitivity to effort costs ^25^. Against this explanation, is that lower effort decision strategies such as preservation were less likely in our apathy patients^88^. We would argue that the observed reduction in the proportion of perseverative choices, is not consistent with a strategy minimising the costs of cognitive control or an indifference to allocate effort or engage in the task.

### Conclusions and future predictions

Allowing for these limitations, our results are in agreement with a view that apathy in PD is unlikely to arise from a single loss of brain circuit function^19, 25, 89^. Conceivably, the first step towards symptomatic manifestation of apathy is a loss of precision in stimulus value encoding ^90^, related to loss of prefrontal dopaminergic projections ^63, 91^. We predict that the expression of demotivated behaviour in PD-apathy arises from secondary loss of compensation using neural decision circuits that can overcome limitations in value encoding. Obvious candidates include noradrenergic systems which are proposed to regulate opponency of the DMN and SN^92^. Noradrenergic reuptake inhibitors are also actively considered as therapy for PD-apathy ^41^ . Serotonin denervation in the dACC correlates with the severity of PD-apathy^32^ and reinnervation in this area reverses apathy^89^. As serotonin promotes choice persistence^34^, upregulation of serotonergic systems could be an explanation for how our non-Apathy group could continue to exploit, despite impaired encoding of a decision outcomes. A ”double-hit” phenomenon, would be consistent with recent longitudinal imaging by Morris et. al., (2023) where loss of functional connectivity between dACC and ventral striatum in non-apathetic PD patients, proceeded the clinical expression of a demotivated state ^93^.

Combining computational modelling of behaviour with neuroimaging of specific neuro-modulatory circuits may answer these predictions. New treatment targets identified with this approach should aim to augment neural circuit compensation identified in this study and protect from the manifestation of apathy in PD. Targeted neuromodulation of regions within this network, could represent a treatment intervention worthy of future investigation.

## Supporting information

Supplementary methods and results

## Acknowledgements

We would like to thank the patients and healthy controls for participating in this study. Thank you to Professor Jens Peters and Prof Nathaniel Daw providing the restless bandit reward payouts used in the study.

## Funding

This work was supported by a Tenovus Scotland research award to T.P.G.

## Competing Interests

The authors report no competing interests.

## Notes

https://osf.io/h3e5s/

### Competing Interest Statement

The authors have declared no competing interest.

