## Supplementary methods and results for "Impaired value-based decision-making in Parkinson’s Disease Apathy"

*^8^SINAPSE www.sinapse.ac.uk*

|  | PD Apathy  LARS>-22 | PD No Apathy  LARS<-22 | Healthy Controls | Control vs Parkinsons | PD Apathy v PD no Apathy |
| --- | --- | --- | --- | --- | --- |
| Participant numbers | 25 | 28 | 22 | n/a | n/a |
| LARS Intellectual Curiosity | -1.45 ± 1.5 | -2.50 ± 1.8 | -3.20 ± 0.5 | **p=0.002** | **p=0.029** |
| LARS Emotion | -1.91 ± 1.9 | -2.95 ± 1.03 | -2.48 ± 1.57 | p=0.811 | **p=0.018** |
| LARS Action Initiation | -1.38 ± 1.42 | -3.09 ± 1.50 | -2.32 ± 1.68 | **p=0.003** | **p<0.001** |
| LARS  Self Awareness | -2.57 ± 1.41 | -3.11 ± 1.34 | -2.86 ± 1.39 | p=0.167 | p=0.342 |

**Supplementary Table 1 Lille Apathy Rating scale Factorial sub-scores.** Values expressed as mean ± standard deviation. P-values in bold, unpaired two-tailed T-test.

**Supplementary Methods**

*Computational modelling*

The probability *P* of choosing a particular bandit, *i* , on trial, *t*, was determined by passing the estimated payout value, $\hat{u}_{i,t}^{pre}$ through a SoftMax function. This process was governed by four different choice rules. *Choice rule 1* (SM) utilized a temperature parameter, *β*, to weigh choices based on their estimated value, thus balancing exploitation and exploration.

*Choice rule 1(SM):-*

$$P_{i,t}=\frac{exp(\beta\hat{u}_{i,t}^{pre})}{\sum_{j} exp(\beta\hat{u}_{j,t}^{pre})}$$

The inverse temperature parameter, $\beta$, weights the sensitivity of the choices by their estimated value. Accordingly, this parameter proportionally scales the degree of exploitation versus exploration, as high values of $\beta$ drives a greedy choice policy that favours bandits with the highest expected value ($\hat{u}_{i,t}^{pre}$).

Choice rule 2 includes an additional term which weights choices by the estimated precision (variance) of each bandit’s value, $\hat{\sigma}_{i,t}^{pre}$. This is done by using an exploration bonus, $\varphi$, which augments uncertainty driven directed exploration:

*Choice rule 2(SME):-*

$$P_{i,t}=\frac{exp(\beta[\hat{u}_{i,t}^{pre}+\varphi\hat{\sigma}_{i,t}^{pre} ])}{\sum_{j} exp(\beta\hat{[u}_{j,t}^{pre}+{\varphi\hat{\sigma}}_{j,t}^{pre}])}$$

Given our *a priori* hypothesis that a perseverative strategy might reflect heighted effort sensitivity in apathy^1,2^, we also included two further choice rules, which influenced choice repetition by weighting each bandit’s choice probability in the model by a preservation bonus, $\rho$ , which is added to the chosen actions probability of being chosen, if two more consecutive trials are repeated, *Ict*−1=*i*.

In both the SMP and SMEP choice rule, the variable represented by *I* , is either 0 or 1 depending upon whether the chosen bandit on trial t is the same as that on the previous trial .

*Choice rule 3 (SMP):-*

$$P_{i,t}=\frac{exp(\beta[\hat{u}_{i,t}^{pre}+ I_{c_{t-1=i}}\rho])}{\sum_{j} exp(\beta[\hat{u}_{j,t}^{pre}+I_{c_{t-1=j}}\rho])}$$

A fourth choice rule included both the exploration and perseveration bonuses**;**

*Choice rule 4 (SMEP):-*

$$P_{i,t}=\frac{exp(\beta[\hat{u}_{i,t}^{pre}+\varphi\hat{\sigma}_{i,t}^{pre}+ I_{c_{t-1=i}}\rho])}{\sum_{j} exp(\beta[\hat{u}_{j,t}^{pre}+{\varphi\hat{\sigma}}_{j,t}^{pre}+I_{c_{t-1=j}}\rho])}$$

In both the SMP and SMEP choice rule, the variable represented by $I$ , is either 0 or 1 depending upon whether the chosen bandit on trial $t$ is the same as that on the previous trial $c_{t-1}$.

*Bayesian Learner*

The Bayesian learner implements the Kalman filter ^3^ to keep track of the estimated value $\hat{u}_{i,t}^{2pre}$ and uncertainty $\hat{\sigma}_{i,t}^{2pre}$ of each bandit $i$ on trial $t$. We initialise these beliefs as $\hat{u}_{all,1}^{2pre}=50$ and $\hat{\sigma}_{all,1}^{2pre}=4$ v ^4^. After a bandit is chosen $c_{t}$the reward for that bandit $i$ is obtained $r_{t}.$ The prediction error $\delta_{t}$is

$$\delta_{t}= r_{t}-\hat{u}_{c_{t},t}^{pre}$$

The Kalman filter, has a variable learning rate (the Kalman gain), where the confidence in the measurement makes a dramatic difference to the amount learned from it. The estimated variance of the given action $\hat{\sigma}_{c_{t},t}^{2pre}$ and the observation variance $\hat{\sigma}_{o}^{2}$ which represents the uncertainty in all observations. $\hat{\sigma}_{o}=4$ for the purpose of fitting due to model degeneration if freed, a problem previously encountered ^4,5^. The Kalman gain $\kappa_{t}$ is given as:

$$\kappa_{t}= \frac{\hat{\sigma}_{c_{t},t}^{2pre}}{\hat{\sigma}_{c_{t},t}^{2pre}+\hat{\sigma}_{o}^{2}}$$

The Kalman gain naturally increases, such that options that have high uncertainty can be quickly estimated, but options that are well established are fine-tuned, with only a small amount learned from trial to trial. The estimated value of the bandit $\hat{u}_{i,t}^{2post}$and the associated variance $\hat{\sigma}_{c_{t},t}^{2post}$ can then be calculated as:

$\hat{u}_{c_{t},t}^{post}= \hat{u}_{c_{t},t}^{pre}+ \kappa_{t}\delta_{t}$	$\hat{\sigma}_{c_{t},t}^{2post}=(1- \kappa_{t}) \hat{\sigma}_{c_{t},t}^{pre}$.

For bandits that are not chosen on any one trial, the prior mean and variance remains unchanged within a trial. However, the prior distributions are updated for all bandits *in between* trials based upon the subjects belief about the gaussian random walk so that;

$\hat{u}_{i,t+1}^{pre}= \hat{u}_{i,t}^{post}+\left( 1-\hat{\lambda} \right)\hat{\vartheta}$ , $\hat{\sigma}_{i,t+1}^{pre}=$ $\lambda^{2}\hat{\sigma}_{i,t}^{post}+ \hat{\sigma}_{d}^{2}$.

Where $\hat{\lambda}$ , $\hat{\vartheta},$ $\hat{\sigma}_{d}$ are constants representing the decay parameter, decay centre and the diffusion variances and were fixed at values of 0.98, 50, and 2.8 for each respective parameter. We used the same values as Chakroun et. al, (2020^4^) which are derived from the actual parameters that govern the gaussian walk the bandits.

*Delta rule*

The delta learning rule has a fixed learning rate,$\alpha$ ,and there is no variance tracking for the uncertainty of the valuation. After a bandit is chosen $c_{t}$the reward for that bandit $i$ is obtained $r_{t}.$ The prediction error $\delta_{t}$is derived in the same way and the prediction of the bandit’s value $\hat{u}_{c_{t},t}^{pre}$ can be updated as such:

$$\delta_{t}= r_{t}-\hat{u}_{c_{t},t}^{pre}, \hat{u}_{c_{t},t+1}^{pre}= \hat{u}_{c_{t},t}^{pre}+ \alpha\delta_{t}$$

There is no decay in this model between trials, the estimated value for a bandit is only updated when that bandit is selected. In the absence of an equivalent estimate of uncertainty $\hat{\sigma}_{c_{t},t}^{pre}$ in the delta learner, we modified choice rules 2 and 4 (SME, SMEP) so that the exploration bonus $\varphi$ was modified by the how long ago a bandit $i$ was chosen $t_{i}$;

*Choice rule 2(SME):*

$$P_{i,t}=\frac{exp(\beta[\hat{u}_{i,t}^{pre}+\varphi(t- t_{i}) ])}{\sum_{j} exp(\beta\hat{[u}_{j,t}^{pre}+\varphi(t- t_{i})])}$$

*Choice rule 4 (SMEP):*

$$P_{i,t}=\frac{exp(\beta[\hat{u}_{i,t}^{pre}+\varphi(t- t_{i})+ I_{c_{t-1=i}}\rho])}{\sum_{j} exp(\beta[\hat{u}_{j,t}^{pre}+\varphi(t- t_{i})+I_{c_{t-1=j}}\rho])}$$

*Model Fitting*

Posterior parameter distributions were estimated for each subject for each of the free parameters specific to the learning ($\alpha$ in the Delta learning rule) and choice rules for each model variant ($\beta, \varphi$, $\rho$*).*

Parameter estimates were derived using Hierarchical Bayesian modelling within Stan (Version 2.17.0; Stan Development Team, 2017), operating in Python version 13. 1.10. Sampling was performed with four chains, each chain running after a warmup period of 5000 iterations for a total of 20000 samples. The prior for each group-level mean was uniformly distributed. For each group-level standard deviation, a half-Cauchy distribution with location parameter 0 and scale parameter 1 was used as a weakly informative prior ^6^. Priors for all subject-level parameters were normally distributed with a parameter-specific mean and standard deviation.

Group level posterior distributions for the free parameters were estimated separately for each patient group (PD-Apathy, PD-Non-apathy and HC). Posterior means and highest density intervals (HDI) were calculated using “bayestestR2 and ‘tidybayes’ packages in R (version 4.1.1 http://www.r-project.org)^7,8^.

*fMRI methods*

For each participant, functional whole-brain images were acquired using a 3T Siemens Prisma Fit scanner using an echo planar imaging sequence with the following parameters: angle = 90°, field of view = 224 mm, matrix = 64 × 64, 37 slices, voxel size 3.5 × 3.5 × 3.5 mm.

Preprocessing was performed in SPM12 (<http://www.fil.ion.ucl.ac.uk/spm>). During this phase, images were realigned, co-registered and normalised to the SPM12 Montreal Neurological Institute (MNI) echo planar imaging template, then smoothed using a 6 mm full-width at half-maximum (FWHM) Gaussian kernel.

The first level General Linear model (GLM) analysis included two time points: the trial onset when the four “bandits” were displayed and the outcome (payout) presentation time. To create regressors for the GLM time points, we convolved these event onsets with the canonical haemodynamic response function (HRF).

For the main GLM, first level contrast images were created for each subject with four session related constants for the five regressors of interest, in the following order: 1) trial onset, 2) outcome onset, 3) choice type, 4) prediction error and 5) outcome value. The trial onset regressor was parametrically modulated by the type of choice as [explore = 1, exploit = 0]. The outcome time was modulated by the outcome value and model prediction error. Orthogonalization between regressors was turned off, as previous work has shown that many of these regressors correlate strongly and could otherwise cancel each other out [44]. Low-frequency noise was removed by employing a temporal high-pass filter with a cut-off frequency of 1/128 Hz, and a first order autoregressive model AR(1) was used to remove serial correlations.

**Supplementary Results**

*In scanner behavioural performance*

The performance trends observed during the Out-of-Scanner session remained consistent in the In-Scanner session (Supplementary Figure 1). We observed the same impairment in best bandit choice in the PD-Apathy group (Best bandit choice: PD-Apathy = 0.5±0.05, PD-noApathy = 0.64±0.04, HC 0.65±0.05, Main effect of Group F(2,279) = 3.96, p= 0.02) and correlation with apathy severity (rho = -0.46, n = 39, p= 0.003). There was also no difference in decision time between groups (F(2,279) = 0.06, = 0.93 or proportion of missed trials F(2,279) = 1.36, = 0.26).

*Model posterior predictive checks*

Replicating previous studies in healthy controls performing the restless bandit task ^4^,^9^ the overall winning model was the Bayes-SMEP (Supplementary Figure 2A). Consistent with this capturing the decisions making in the task robustly, model parameters could still be successfully recovered from synthetic choice data generated from simulated choices derived from individual subjects' model parameters (Supplementary Figure B-D). We found further support of a good model fit of the experimental data by showing that the simulated data choices overlapped with the experimental group data (Supplementary Figure 3 A-B, D-E). Furthermore, the relationship between simulated choices from the individual subjects LARS score could also be reproduced from the simulated data (Supplementary Figure 3 C&F).

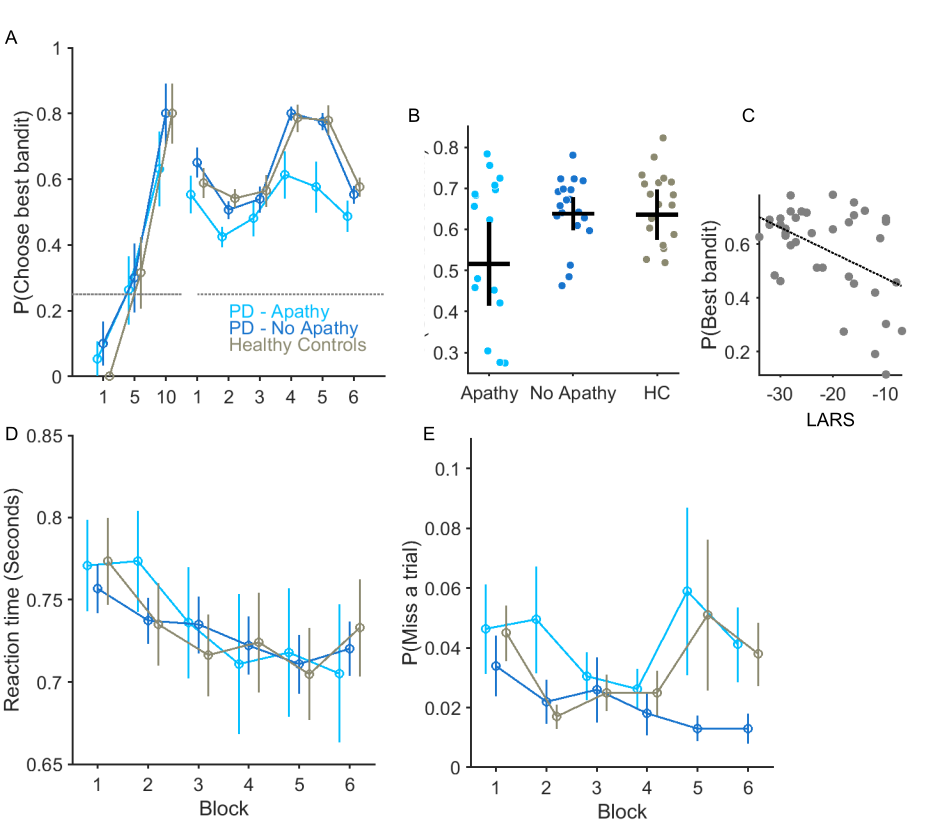

**Supplementary Figure 1 Bandit performance and relationship with Apathy Severity (in Scanner).** The average probability of choosing the bandit with the highest payout, P(Choose best bandit) is plotted the three groups of participants. (**A**) The increase in average values of best bandit choice plotted for in trials 1-10 confirm learning in all three groups from an initial random choice to above chance levels( Best bandit choice performance across six 50 trial bins of the task was reduced in the PD-Apathy group. Vertical lines S.E.M. In **(B**) each circle represents average best bandit choice probability across the task for an individual subject with the horizontal and vertical bar represents the group mean and 95% confidence limits. (**C**) Apathy severity, measured by increasing LARS score (more positive values represent higher levels of apathy) correlated with individuals ability to choice the best bandit (r = -0.46, p= 0.003). Each group’s average reaction time over the 6 task bins in (**D**) and the likelihood of not making a response (**E**) (missed trial) was not affected by apathy.

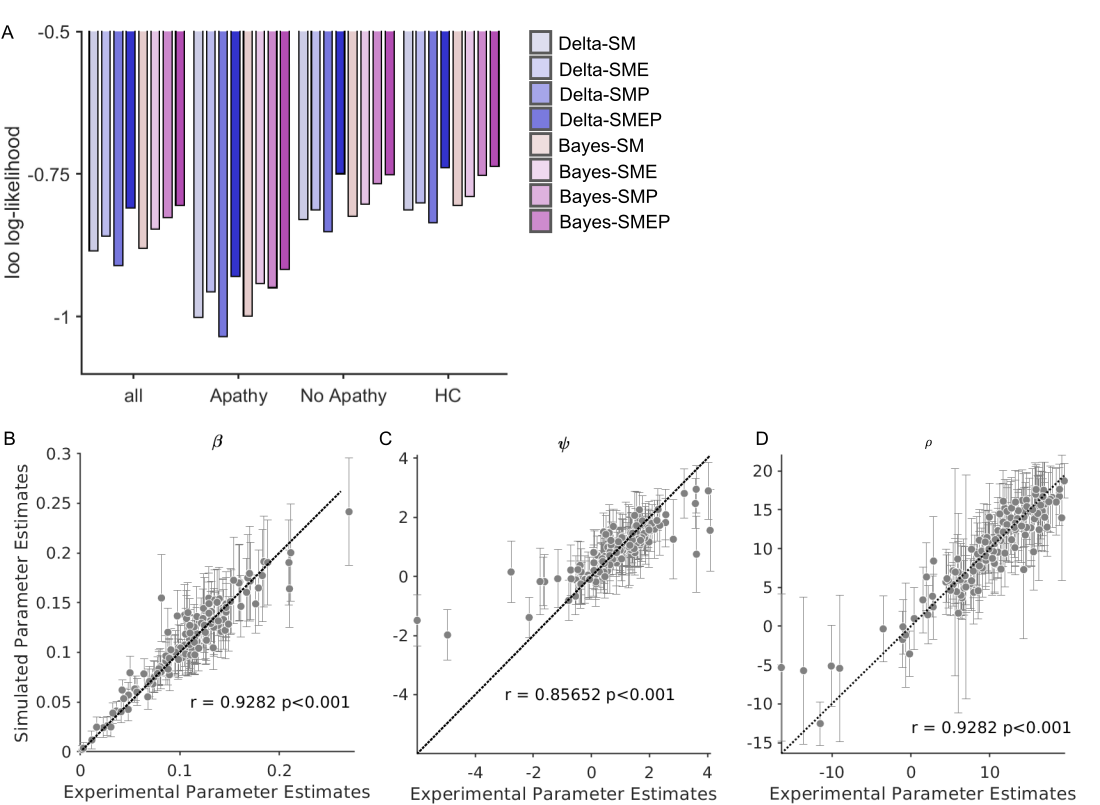

**Supplementary Figure 2 Model Fit and Parameter recovery. (A**) Loo-log likelihood scores combined from all modelling fitting for each model variant (two learning rules, Bayes & Delta, four choice rules: SM: Softmax, SME: Softmax with exploration bonus, SMP: Softmax with preservation bonus, SMEP: Softmax with exploration and perseveration bonus. More positive values of loo-log like values indicate better model fit. Ten simulated datasets were generated per participant for all three groups. Refitting these simulations recovered the same individual parameter estimates from the experimental choices for the three free parameters in the winning Bayes-SMEP model. Each circle represents the parameter value from the experimental choice fitting plotted against the average parameter estimate from the 10 simulations. Error bars are 95% confidence limits (**B**) reward sensitivity parameter $\beta$ (**C**) exploration bonus $\varphi$ and perseveration bonus $\rho$ (**D**).

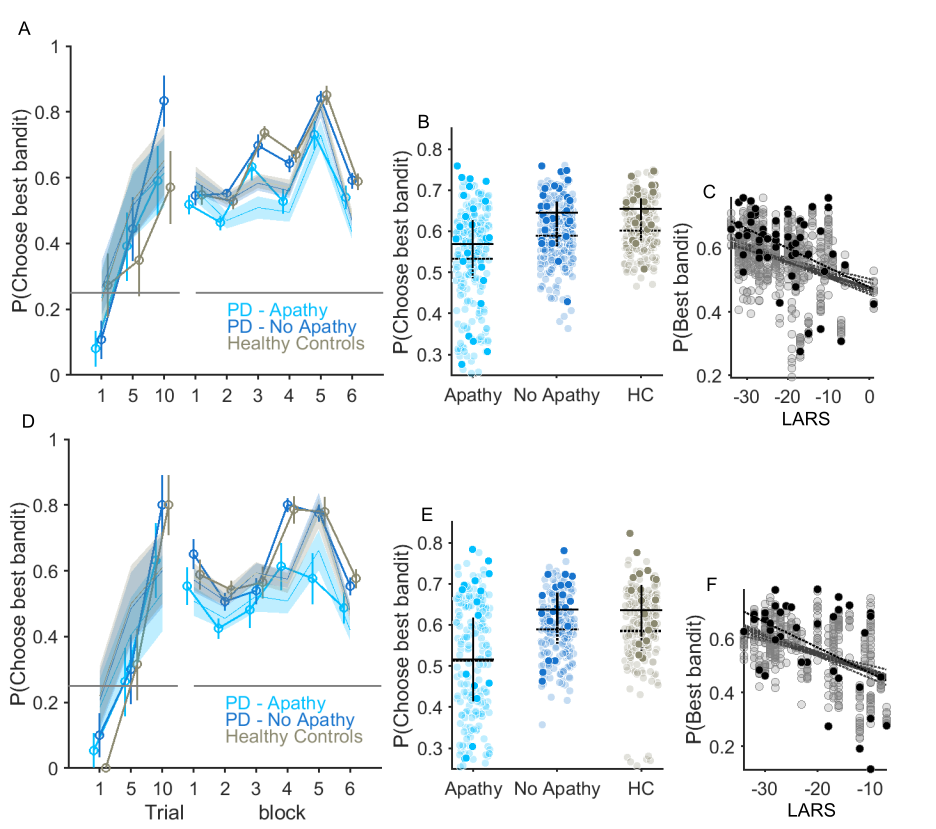

**Supplementary Figure 3 Overlap between simulated and observed experimental data**. (**A-C**) Restless bandit performance from “out-of-scanner” sessions. (**D-F**) Experimental performance and model simulations from In-Scanner session. Overlayed onto the experimental performance for each group is the average best bandit choice probability across six-50 trial bins from the Bayes-SMEP model simulations (10 simulations used for parameter recovery). The between simulation error is represented by the 95% confidence limit in the shaded colour coded regions for each group in (**A**). In (**B**) each dark circle represents the experimental average best bandit choice probability across the whole task with the simulated equivalent overlayed the lighter coloured circles. Solid horizontal cross hairs illustrate the group average experimental performance, vertical the 95% confidence limits. The average simulated performance and averaged between simulation error is represented by the horizontal and vertical dashed lines. (**C**) The dark circles and dark line of best fit from correlating Apathy severity (LARS) with experimental task performance are plotted overlayed with the same analysis applied to the 10 stimulations for the PD-Apathy and PD-No Apathy groups (i.e. the same as that presented in Figure 2C). A significant negative correlation (p<0.05) was between LARS score and task performance was identified from 9 of the 10 simulations in the out-of-scanner and from all of the In-scanner simulated data (**F**).

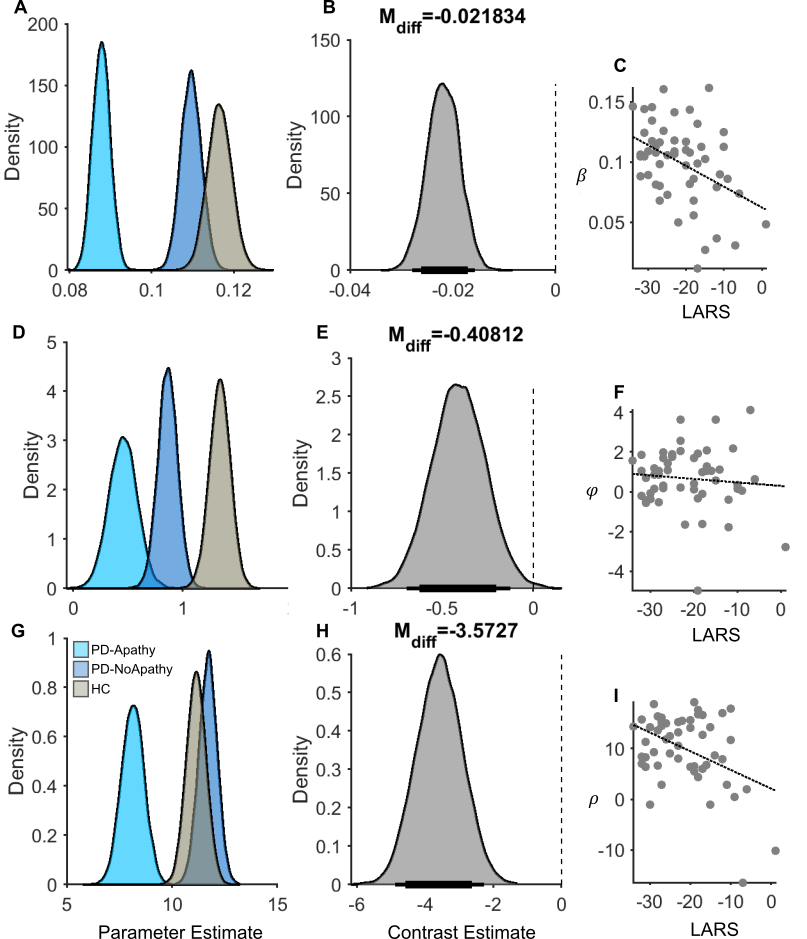

**Supplementary Figure 4 Posterior distributions and contrast estimates for the Bayes-SMEP model.** Group level parameter estimates for the reward sensitivity parameter $\beta$ **(A),**exploration (**D**)$\varphi$, and perseveration bonuses $\rho$ (**G**). Posterior distributions of differences in the mean (M_diff_) PD-Apathy minus PD-NoApathy for the corresponding parameter (**B-H**). Thick and thin horizontal bars below the distributions represent the 85% and 95% highest density intervals, respectively. The vertical dotted line highlights the zero point of no difference in the parameter means. Correlation between the individual apathy score in the patients (LARS) with their model parameter estimate. Significant correlations were identified with both $\beta$ the r = -0.41, p=0.002 (**C**) and $\rho$ (**I**) r = -0.41, p=002 but not $\varphi$ (**F**).

|  | **Model Parameter** | | | | | | | | |
| --- | --- | --- | --- | --- | --- | --- | --- | --- | --- |
| **Group** | $\beta$ (reward sensitivity) | | | $\varphi$ (exploration bonus) | | | $\rho$ (perseveration bonus) | | |
|  | **Mean** | **95% HDI** | | **Mean** | **95% HDI** | | **Mean** | **95% HDI** | |
| **PD-Apathy** | 0.08 | 0.08 | 0.09 | 0.45 | 0.30 | 0.70 | 8.13 | 7.13 | 9.20 |
| **PD-NoApathy** | 0.11 | 0.10 | 0.11 | 0.86 | 0.68 | 1.03 | 11.70 | 10.85 | 12.56 |
| **HC** | 0.115 | 0.11 | 0.12 | 1.33 | 1.14 | 1.51 | 11.15 | 10.25 | 12.05 |
| **PD Apathy v NoApathy** | -0.02* | -0.028 | -0.015 | -0.02* | -0.028 | -0.015 | -3.57* | -4.88 | -2.27 |
| **Correlation with LARS** | -0.41, p = 0.002 | | | 0.0048, p=0.51 | | | -0.41, p=0.002 | | |

**Supplementary Table 2 Summary of Posterior distributions and contrast estimates of the Bayes-SMEP model parameters**. *Denotes Significant differences in the posterior distributions of the parameter estimate at 95% confidence limit.

*In-Scanner Model Fitting*

In the analysis of choices made in the scanner, we observed a consistent relationship between the severity of apathy (as indexed by LARS Score) and a propensity for heightened exploration and reduced exploitation. This was indicated by P(RE) versus LARS with rho(38) = 0.38, p=0.01, and P(Exploit) versus LARS with rho(38) = -0.38, p=0.01.

However, we did not replicate the main effect of group on average choices. For instance, the proportion of exploitative choices, P(Exploit), were PD - Apathy = 0.62±0.06, PD-noApathy = 0.73±0.02, HC = 0.74±0.03, with a Main effect of Group F(2,279) = 2.51, p=0.09. Similarly, the proportions of random exploration, P(RE), were PD-Apathy = 0.24±0.04, PD-noApathy = 0.16±0.020, HC = 0.15±0.03, with a Main effect of Group F(2,279) = 2.08, p=0.13. Despite this, we did confirm a significant reduction in the inverse temperature parameter *β* (M_diff_ = -0.02 [-0.03, -0.011]), and exploration bonus *φ* (M_diff_ = -0.60[-0. 96, -0.23]) in the apathy group. There were no significant differences in levels of perseveration among groups (P(Stay): PD - Apathy = 0.58±0.05, PD-noApathy = 0.66±0.04, HC = 0.69±0.05; Main effect of Group F(2,279) = 1.11, p=0.33), and there was also no notable difference in the estimate for the perseveration bonus, *ρ* (Supplementary Figure 5 & Supplementary Table 4).

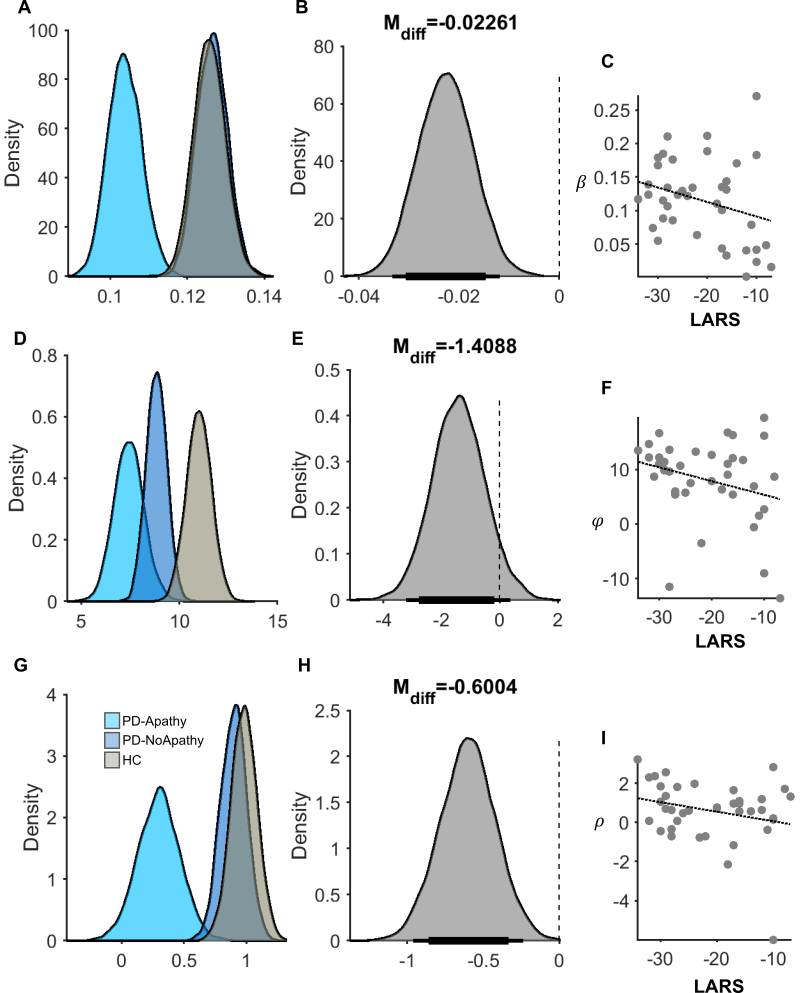

**Supplementary Figure 5 Posterior distributions and contrast estimates for the Bayes-SMEP model – In Scanner.** Group level parameter estimates for the reward sensitivity parameter $\beta$ **(A),**exploration (**D**)$\varphi$, and perseveration bonuses $\rho$ (**G**). Posterior distributions of differences in the mean (M_diff_) PD-Apathy minus PD-NoApathy for the corresponding parameter (**B-H**). Thick and thin horizontal bars below the distributions represent the 85% and 95% highest density intervals, respectively. The vertical dotted line highlights the zero point of no difference in the parameter means. Correlation between the individual apathy score in the patients (LARS) with their model parameter estimate. No significant correlations were identified with the apathy severity and the model parameters $\beta$ (**C**) ,$\rho$ (**I**) or $\varphi$ (**F**).

|  | **Model Parameter** | | | | | | | | |
| --- | --- | --- | --- | --- | --- | --- | --- | --- | --- |
| **Group** | $\beta$ (reward sensitivity) | | | $\varphi$ (exploration bonus) | | | $\rho$ (perseveration bonus) | | |
|  | **Mean** | **95% HDI** | | **Mean** | **95% HDI** | | **Mean** | **95% HDI** | |
| **PD-Apathy** | 0.1 | 0.09 | 0.11 | 0.29 | -0.01 | 0.64 | 7.42 | 5.99 | 8.97 |
| **PD-NoApathy** | 0.12 | 0.11 | 0.13 | 0.89 | 0.69 | 1.09 | 8.83 | 7.86 | 9.89 |
| **HC** | 0.12 | 0.11 | 0.13 | 0.98 | 0.78 | 1.29 | 10.99 | 9.76 | 12.24 |
| **PD Apathy v NoApathy** | -0.02* | -0.03 | -0.011 | -0.60* | -0.96 | -0.23 | -1.41 | -3.21 | 0.37 |
| **Correlation with LARS** | -0.28 p = 0.07 | | | 0.24, p=0.12 | | | 0.27, p=0.09 | | |

**Supplementary Table 4 Summary of Posterior distributions and contrast estimates of the Bayes-SMEP model parameters – In Scanner**. *Denotes Significant differences in the posterior distributions of the parameter estimate

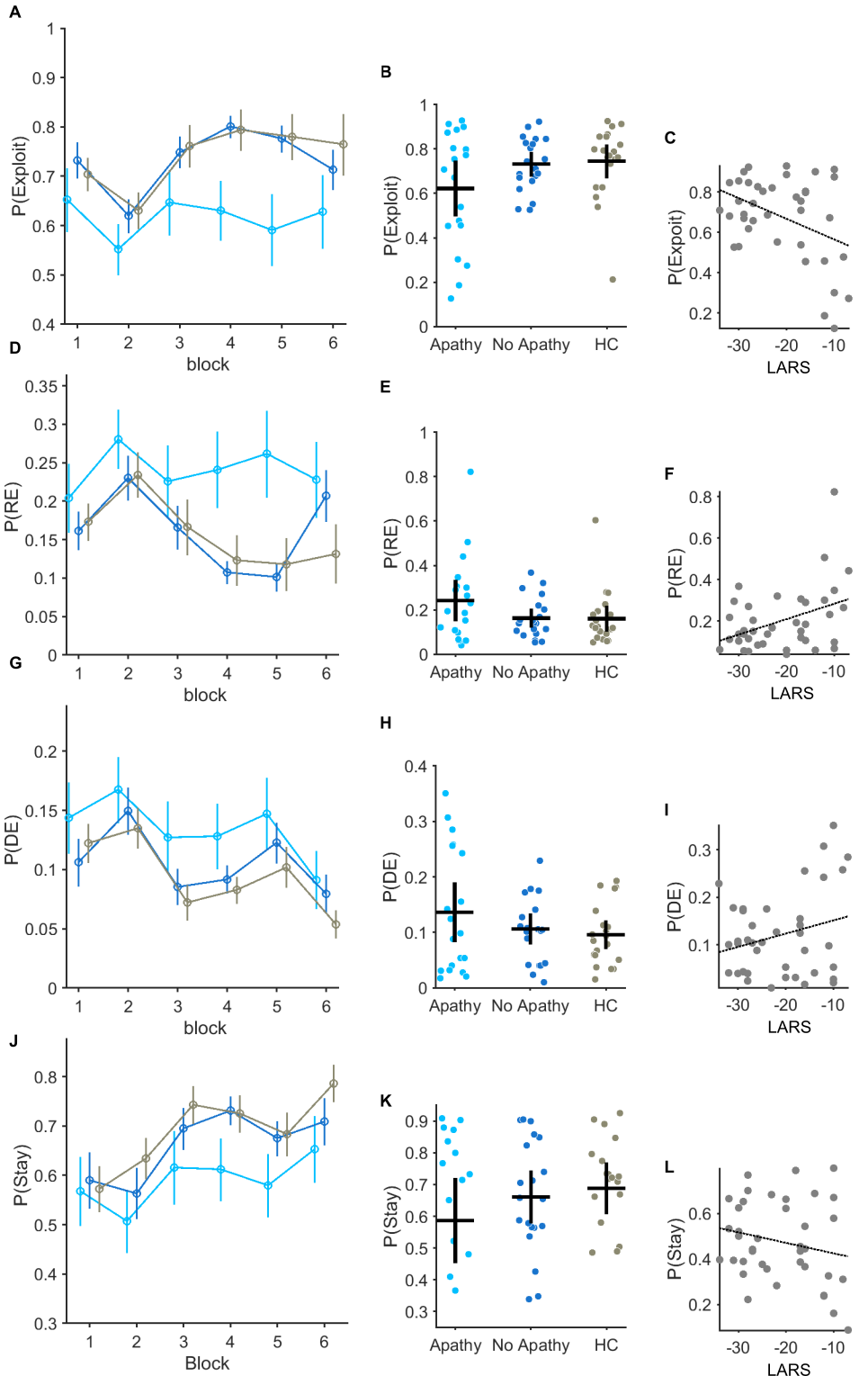

**Supplementary Figure 6 Decision types between groups across the task and relationship with Apathy severity - In Scanner.** **(A-C)** Probability of making an exploit P(Exploit) choice plotted across six-50 trial bins for each group tested (**A**). Error bars represent S.E.M. **(B)** individual P(Exploit) across the whole task is represented by each circle. Group average and S.E.M is illustrated by the vertical and horizontal bars. Correlation between P(Exploit) and Apathy severity (increasing LARS Score) in (**C**) r = -0.38, p=0.01. The same analysis applied to the probability of making a random exploratory P(Explore) choice **(D-E)** and relationship with individual apathy severity (F) r = 0.38, p=01. P(DE) is the probability of making a directed exploratory choice (**G-I**) and P(Stay) the same choice on two consecutive trials (**J-K**).

*fMRI activations and between group contrasts*

| **Activations – Outcome** | | | | | | |
| --- | --- | --- | --- | --- | --- | --- |
| **Age matched healthy controls (HC)** | | | | | | |
| **Region** | **MNI coordinates** | | | **Hemisphere** | **Peak**  **T-value** | **Cluster**  **Extent (k)** |
|  | x | y | z |  |  |  |
| Ventromedial prefrontal cortex | -6 | 34 | -8 | L  L  R | 5.60 | 590 |
|  | 4 | 28 | -14 |  | 4.25 |  |
|  | -6 | 24 | -16 |  | 4.23 |  |
| **Parkinson’s Disease with Apathy (PD Apathy)** | | | | | | |
| *No Suprathreshold clusters* | | | | | | |
| **Parkinson’s Disease without Apathy (PD No Apathy)** | | | | | | |
| *No Suprathreshold clusters* | | | | | | |
| **Contrast PD Apathy < PD No Apathy** | | | | | | |
| *No Suprathreshold clusters* | | | | | | |
| **Contrast PD Apathy > PD No Apathy** | | | | | | |
| *No Suprathreshold clusters* | | | | | | |
| **Contrast PD No Apathy < HC** | | | | | | |
| *No Suprathreshold clusters* | | | | | | |
| **Contrast PD Apathy < HC** | | | | | | |
| *No Suprathreshold clusters* | | | | | | |

**Supplementary Table 5 Summary of fMRI activations to the outcome in each group and between group contrasts.** All Clusters reported survive cluster-level FWE correction at *P* < 0.05 with an initial uncorrected cluster-defining threshold of *P* < 0.001 and a minimum cluster size of 10 voxels.

| **Activations – Explore** | | | | | | |
| --- | --- | --- | --- | --- | --- | --- |
| **Parkinson’s Disease with Apathy (PD Apathy)** | | | | | | |
| **Region** | **MNI coordinates** | | | **Hemisphere** | **Peak**  **T-value** | **Cluster**  **Extent (k)** |
|  | x | y | z |  |  |  |
| Calcarine Cortex | -12 | -86 | 16 | L  L  R | 5.40 | 1802 |
|  | -8 | -92 | 6 |  | 5.25 |  |
|  | 12 | -88 | 6 |  | 5.12 |  |
| Fusiform gyrus | 34 | -64 | -14 | R | 5.02 | 590 |
|  | 30 | -74 | -18 |  | 4.01 |  |
|  | 30 | -54 | -14 |  | 3.91 |  |
| Superior Parietal lobule  Intraparietal sulcus | -36 | -54 | 56 | L | 4.77 | 1311 |
|  | 30 | -74 | -18 |  | 4.60 |  |
|  | -18 | -60 | 58 |  | 4.45 |  |
| Insula | -28 | 22 | 0 | L | 4.72 | 310 |
|  | -28 | 26 | 8 |  | 4.24 |  |
|  | -26 | 20 | 10 |  | 4.17 |  |
| Middle Frontal Gyrus (dorsal Prefrontal cortex)  Middle Frontal Gyrus (Premotor cortex) | 42 | 28 | 38 | R | 4.53 | 1415 |
|  | 36 | 0 | 58 |  | 4.40 |  |
|  | 60 | 12 | 38 |  | 4.32 |  |
| Superior frontal gyrus (dorsomedial prefrontal cortex)  preSMA/dorsal anterior cingulate cortex | -4 | 36 | 44 | L | 4.44 | 1732 |
|  | 8 | 24 | 50 | R  R | 4.27 |  |
|  | 10 | 26 | 34 |  | 4.27 |  |
| Supramarginal Gyrus | 48 | -30 | 44 | R | 4.40 | 160 |
| Intraparietal sulcus | 38 | -76 | 26 | R | 4.22 | 291 |
|  | 34 | -70 | 42 |  | 4.04 |  |
|  | 30 | -70 | 50 |  | 3.87 |  |
| Middle Frontal Gyrus (Premotor cortex) | -52 | 12 | 32 | L | 4.15 | 455 |
|  | -38 | 26 | 42 |  | 4.05 |  |
|  | -42 | 22 | 28 |  | 4.02 |  |
| Middle Frontal Gyrus (Frontal Pole) | -40 | 40 | 12 | L | 3.89 | 177 |
|  | -30 | 50 | 18 |  | 3.75 |  |
|  | -50 | 40 | 20 |  | 3.58 |  |
| **Parkinson’s Disease without Apathy (PD No Apathy) *** | | | | | | |
| Middle Frontal Gyrus (Premotor cortex)  Insula | -46 | 4 | 30 | L | 8.64 | 3365 |
|  | -30 | 20 | -10 | L | 7.33 |  |
|  | -40 | 10 | 30 | L | 7.32 |  |
| Calcarine cortex  Bilateral Midbrain/ Thalamus  Middle Frontal Gyrus (dorsal prefrontal cortex/premotor cortex)  Right Frontal pole  Insula  IPS  Cerebellum | -6 | -90 | 0 | L | 8.40 | 38501 |
|  | 10 | -20 | -8 | R | 8.22 |  |
|  | 26 | -54 | -4 | R | 7.98 |  |
| preSMA/  dorsal anterior cingulate cortex | 8 | 10 | 52 | R | 5.5 | 2338 |
|  | -2 | 4 | 54 | L | 5.45 |  |
|  | 6 | 16 | 46 | R | 5.39 |  |
| Primary sensory cortex | -52 | -20 | 18 | L | 5.55 | 286 |
|  | -64 | -16 | 18 | L | 4.84 |  |
| Frontal Pole | -28 | 54 | -6 | L | 5.06 | 357 |
|  | -20 | 52 | -16 | L | 4.72 |  |
|  | -36 | 52 | -4 | L | 4.68 |  |
| Primary sensory cortex | 60 | -14 | 22 | R | 4.66 | 84 |
|  | 54 | -14 | 30 | R | 4.60 |  |
| **Age matched healthy controls (HC)** | | | | | | |
| Calcarine cortex/  Superior parietal lobule Intraparietal sulcus  Pre-post central gyrus | 16 | -88 | 10 | R | 7.57 | 15700 |
|  | -8 | -94 | 8 | L | 6.67 |  |
|  | 10 | -68 | 56 | R | 6.52 |  |
| Middle Frontal Gyrus  (Dorsal Prefrontal cortex  /Premotor cortex) | 46 | 32 | 30 | R | 5.55 | 1045 |
|  | 38 | 8 | 32 | R | 4.29 |  |
|  | 48 | 8 | 39 | R | 4.13 |  |
| Insula | 30 | 24 | -4 | R | 5.50 | 292 |
|  | 40 | 18 | 8 | R | 3.52 |  |
| Middle Frontal Gyrus  (Premotor cortex/Dorsal Prefrontal cortex) | -42 | 8 | 32 | L | 5.50 | 1928 |
|  | -24 | -2 | 46 | L | 5.18 |  |
|  | -46 | 30 | 26 | L | 4.80 |  |
| Insula | -28 | 22 | -4 | L | 5.07 | 407 |
|  | -28 | 20 | 6 | L | 5.01 |  |
|  | -44 | 14 | 2 | L | 3.56 |  |
| preSMA/dACC | -8 | 16 | 46 | L | 5.06 | 520 |
|  | 6 | 22 | 48 | R | 4.10 |  |
| Midbrain  Midbrain  Thalamus | 8 | -24 | -8 | R | 4.61 | 290 |
|  | -6 | -20 | -8 | L | 4.33 |  |
|  | -14 | -14 | -6 | L | 3.34 |  |
| Thalamus | 10 | -12 | 2 | R | 4.07 | 134 |

**Supplementary Table 6 Summary of fMRI activations at the decision time during exploration.** All activations surviving cluster-level FWE correction at *P* < 0.05 with an initial uncorrected cluster-defining threshold of *P* < 0.001 or * P<0.0001 (with k ≥10 voxels) *

| **Activations – Exploit** | | | | | | |
| --- | --- | --- | --- | --- | --- | --- |
| **Parkinson’s Disease with Apathy (PD Apathy)** | | | | | | |
| **Region** | **MNI coordinates** | | | **Hemisphere** | **Peak**  **T-value** | **Cluster**  **Extent (k)** |
|  | x | y | z |  |  |  |
| *No Suprathreshold clusters* | | | | | | |
| **Parkinson’s Disease without Apathy (PD No Apathy)** | | | | | | |
| *No Suprathreshold clusters* | | | | | | |
| **Age matched healthy controls (HC)** | | | | | | |
| Superior Frontal Gyrus | -18 | 52 | 22 | L | 4.63 | 194 |
|  | -10 | 54 | 18 | L | 4.18 |  |
| Ventromedial PFC  subgenual cingulate cortex | -18 | 36 | 4 | L | 4.13 | 389 |
|  | -16 | 30 | -4 | L | 3.93 |  |
|  | -10 | 26 | -12 | L | 3.79 |  |
| **Contrasts - Exploit** | | | | | | |
| **PD Apathy< PD No Apathy** | | | | | | |
| *No Suprathreshold clusters* | | | | | | |
| **PD Apathy>PD No Apathy** | | | | | | |
| *No Suprathreshold clusters* | | | | | | |

**Supplementary Table 7 Summary of fMRI activations at the decision time during exploitation.** All Clusters reported survive cluster-level FWE correction at *P* < 0.05 with an initial uncorrected cluster-defining threshold of *P* < 0.001 and a minimum cluster size of 10 voxels.

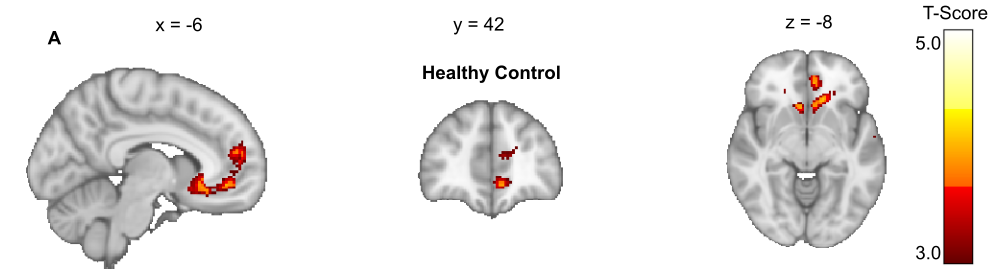

**Supplementary Figure 6 fMRI BOLD signal correlates at the decision time during exploitative choices.** Peak activations in healthy control (HC) group at the point of decision making when an exploitative choice was made activated superior frontal grrus, vmPFC and subgenual cingulate cortex. No significant clustered survived FWE correction in either the PD-Apathy or PD- No-Apathy Groups (supplementary Table 5).

| **Contrasts - Explore** | | | | | | |
| --- | --- | --- | --- | --- | --- | --- |
| **PD Apathy < PD No Apathy** | | | | | | |
| **Region** | **MNI coordinates** | | | **Hemisphere** | **Peak**  **T-value** | **Cluster**  **Extent (k)** |
|  | x | y | z |  |  |  |
| Thalamus/  Midbrain | 14 | -12 | 2 | R | 5.37 | 3824 |
|  | -20 | -14 | -6 | L | 5.25 |  |
| L Lingual Gryus/ Visual association cortex | 26 | -52 | -4 | R | 4.89 | 140 |
|  | 18 | -60 | 4 | R | 3.87 |  |
| Pre and post central gyri | 6 | -32 | 72 | R | 4.67 | 192 |
|  | -6 | -38 | 72 | L | 3.86 |  |
| Cerebellum | -10 | -58 | -44 | L | 4.57 | 229 |
|  | -16 | -52 | -44 |  | 3.77 |  |
|  | -26 | -50 | -46 |  | 3.59 |  |
| Intraparietal sulcus (IPS) | 26 | -66 | 28 | R | 4.46 | 116 |
| Calcarine cortex | 24 | -90 | -8 | R | 4.33 | 236 |
|  | 30 | -86 | 2 | R | 4.04 |  |
|  | 34 | -84 | -14 | R | 4.01 |  |
| cerebellum | 26 | -52 | -42 | R | 4.33 | 324 |
|  | 14 | -48 | -44 | R | 4.04 |  |
|  | 24 | -58 | -48 | R | 4.01 |  |
| **PD Apathy > PD No Apathy** | | | | | | |
| *No Suprathreshold clusters* | | | | | | |
| **PD Apathy < HC** | | | | | | |
| *No Suprathreshold clusters* | | | | | | |
| **PD Apathy > HC** | | | | | | |
| Frontal Pole | -10 | 54 | 18 | L | 4.67 | 252 |
|  | -18 | 52 | 22 | L | 4.53 |  |
|  | 6 | 52 | 26 | R | 3.57 |  |
| **PD No Apathy < HC** | | | | | | |
| *No Suprathreshold clusters* | | | | | | |
| **PD No Apathy > HC** | | | | | | |
| Pre/post central gyrus | 0 | -30 | 68 |  | 5.52 | 214 |
|  | -8 | -32 | 60 | L | 3.44 |  |
| Calcarine cortex | 32 | -86 | -10 | R | 4.96 | 412 |
|  | 24 | -86 | -30 | R | 4.40 |  |
|  | 32 | -82 | -20 | R | 4.39 |  |
| Supramarginal Gyrus  Primary Motor  L superior Tempral gyrus | -54 | -26 | 16 | L | 4.10 | 372 |
|  | -62 | -12 | 16 | L | 3.85 |  |
|  | -62 | -42 | 14 | L | 3.61 |  |
| Inferior Frontal Gyrus | -32 | 26 | -14 | L | 4.35 | 145 |
|  | -42 | 20 | -20 | L | 3.52 |  |
| cerebellum | -26 | -52 | -42 | L | 4.28 | 415 |
|  | -12 | -58 | -46 | L | 4.15 |  |
|  | -20 | -38 | -46 | L | 3.72 |  |
| Frontal Pole | 8 | 54 | 12 | R | 4.20 | 246 |
|  | -18 | 52 | 22 | L | 4.15 |  |
|  | 14 | 60 | 16 | R | 3.77 |  |

**Supplementary Table 8 Summary of fMRI activation contrasts at the decision time during exploration.** All Clusters reported survive cluster-level FWE correction at *P* < 0.05 with an initial uncorrected cluster-defining threshold of *P* < 0.001 and a minimum cluster size of 10 voxels.
